# Efficient coding of natural scenes improves neural system identification

**DOI:** 10.1101/2022.01.10.475663

**Authors:** Yongrong Qiu, David A. Klindt, Klaudia P. Szatko, Dominic Gonschorek, Larissa Hoefling, Timm Schubert, Laura Busse, Matthias Bethge, Thomas Euler

## Abstract

*Neural system identification* aims at learning the response function of neurons to arbitrary stimuli using experimentally recorded data, but typically does not leverage normative principles such as efficient coding of natural environments. Visual systems, however, have evolved to efficiently process input from the natural environment. Here, we present a normative network regularization for system identification models by incorporating, as a regularizer, the *efficient coding* hypothesis, which states that neural response properties of sensory representations are strongly shaped by the need to preserve most of the stimulus information with limited resources. Using this approach, we explored if a system identification model can be improved by sharing its convolutional filters with those of an autoencoder which aims to efficiently encode natural stimuli. To this end, we built a hybrid model to predict the responses of retinal neurons to noise stimuli. This approach did not only yield a higher performance than the “stand-alone” system identification model, it also produced more biologically-plausible filters. We found these results to be consistent for retinal responses to different stimuli and across model architectures. Moreover, our normatively regularized model performed particularly well in predicting responses of direction-of-motion sensitive retinal neurons. In summary, our results support the hypothesis that efficiently encoding environmental inputs can improve system identification models of early visual processing.

## Introduction

In the past years, advances in experimental techniques enabled detailed, large-scale measurements of activity at many levels of sensory processing (1). As a consequence, *neural system identification (SI)* approaches have flourished (Fig. 1a top). They empirically fit the stimulus-response (transfer) function of neurons based on experimentally recorded data (2–4). A classic example is the generalized linear model (GLM, (2, 5)), which consists of a linear filter as a first order approximation of a neuron’s response function (i.e., its receptive field; (6)), followed by a point-wise nonlinear function for the neuron’s output. To account for additional non-linearities (e.g., (7, 8)), several extensions, such as linear-nonlinear cascades (9, 10), have been proposed. More recently, deep neural network-based SI approaches inspired by the hierarchical processing along the visual pathway (11, 12) have been developed (reviewed in (13–17)). While SI methods became particularly successful in predicting responses of visual neurons (18–22), they often require large amounts of training data and, more critically, do rarely consider adaptions to the natural environment.

**Fig. 1.**
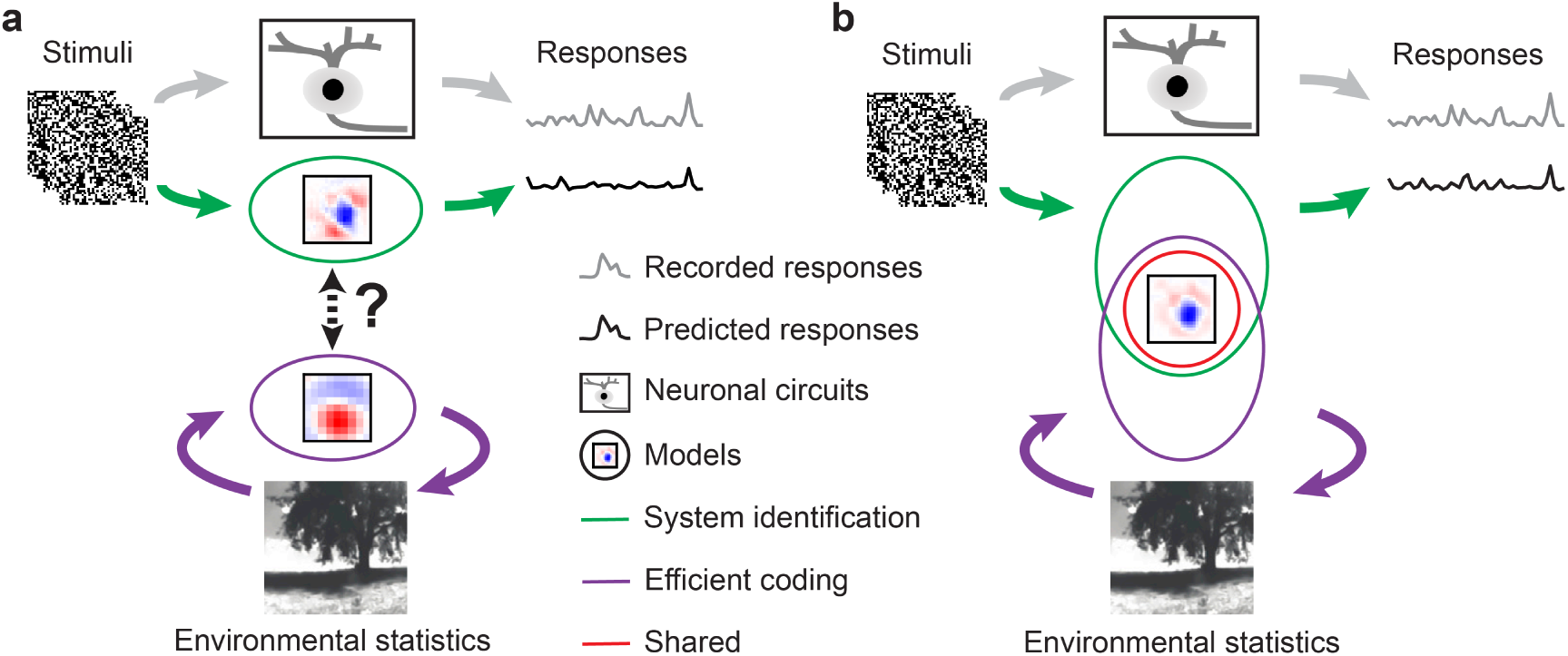
Illustration of our hybrid model combining SI and EC. **a**. Illustration of two common approaches to studying visual systems: system identification, symbolized by the green-labeled branch, aims at predicting responses of neuronal circuits (black rectangle) to specific stimuli, whereas efficient coding (purple-labeled branch) seeks working out principles of the visual system based on environmental statistics. As these two approaches are rarely combined in a single modeling framework, their potential synergies remain largely unexplored. **b**. Our hybrid modeling approach combines system identification (green) and efficient coding (purple) in a single model with shared filters (red circle) to predict neural responses to arbitrary visual stimuli.

However, like other senses, vision has evolved to promote a species’ survival in its natural environment (23), driving visual circuits to efficiently represent information under a number of constraints, including metabolic limits and space restrictions (24, 25). As a consequence, the visual system has adapted to natural statistics, as shown, for example, by the fact that the distribution of orientation preferences of visual neurons mirrors the dominance of cardinal orientations in natural scenes (26–28).

Such adaptations are at the heart of *efficient coding* (EC) approaches (Fig. 1a bottom): They derive algorithmic principles underlying neural systems from the statistical properties of natural stimuli and by incorporating biological constraints (15, 24, 25, 29–31). Here, one popular strategy starts from the assumption that early visual processing serves to decorrelate the redundant signals in natural environments (32, 33). This theory can reproduce feature selectivity, e.g., difference-of-Gaussian (DoG) kernels that have similar receptive field (RF) properties as retinal ganglion cells (RGCs; (34)). Recently, deep neural networks-augmented EC approaches were proposed, such as convolutional autoencoders (35, 36), which are trained to optimally reconstruct inputs in the presence of an information “bottleneck” (i.e., from a constrained latent representation). Such convolutional autoencoders have been shown to yield center-surround spatial RFs with similar properties as those observed in RGCs when encoding either pink (1*/f*) noise or natural scenes (37, 38). Still, a downside of EC is that it is not always straightforward to experimentally measure coding efficiency and feature selectivity predicted by these approaches in neural systems (discussed in (39, 40)) and, hence, the interpretation of EC models with respect to the biological underpinnings remains challenging.

Notably, the intersection between EC and SI has long remained largely unexplored but lately shifted more into focus. In particular, Mlynarski and colleagues recently proposed a theoretical framework incorporating normative theories for statistical inference on simulated or pre-fit neural data (41). Their framework enables conducting rigorous statistical hypothesis tests of coding principles, but has not yet been applied to predicting neural responses to arbitrary stimuli with minimal assumptions. Here, we tested whether the EC hypothesis can serve as a useful inductive bias for learning the response functions of neurons. To do so, we built a hybrid model combining a SI branch with an EC branch, forced the two branches to share filters (Fig. 1b) and asked, if knowledge about natural scene statistics could help predicting retinal responses. To this end, we experimentally recorded Ca^2+^ signals of neurons in the mouse retina while presenting it with visual stimuli and then used these responses to train the SI branch, which aims to predict retinal responses. We used natural movies that we recorded in mouse habitats outdoors to train the EC branch, which aims to represent natural scenes efficiently (38). We found a synergy between neural prediction and natural scene statistics: The hybrid approach did not only have a better predictive performance than a pure SI approach, it also produced more biologically-plausible filters. Our results demonstrate that predicting sensory responses benefits from considering adaptations to the natural environment.

## Results

### Hybrid system identification and efficient coding models

To test if learning an efficient representation of natural input could help predict neuronal responses in the early visual system, we employed *normative regularization*, i.e., statistical regularization that is informed by normative coding principles, such as the idea that sensory systems have evolved to efficiently process natural stimuli. Specifically, we used this strategy to incorporate EC as a regularizer and developed a hybrid model that combines SI-based neural prediction and EC in a single model. The two model branches are linked by shared convolutional filters (Fig. 1b).

The *SI branch* approximates the response functions of recorded neurons to a visual dense noise (see below), and was implemented using a convolutional neural network (CNN) (Fig. 2a). Here, we used an L2 regularization on the convolutional layers to encourage smooth filters (42) and an L1 regularization on the fully connected (FC) layer for sparse readouts ((19); for details, see Methods).

**Fig. 2.**
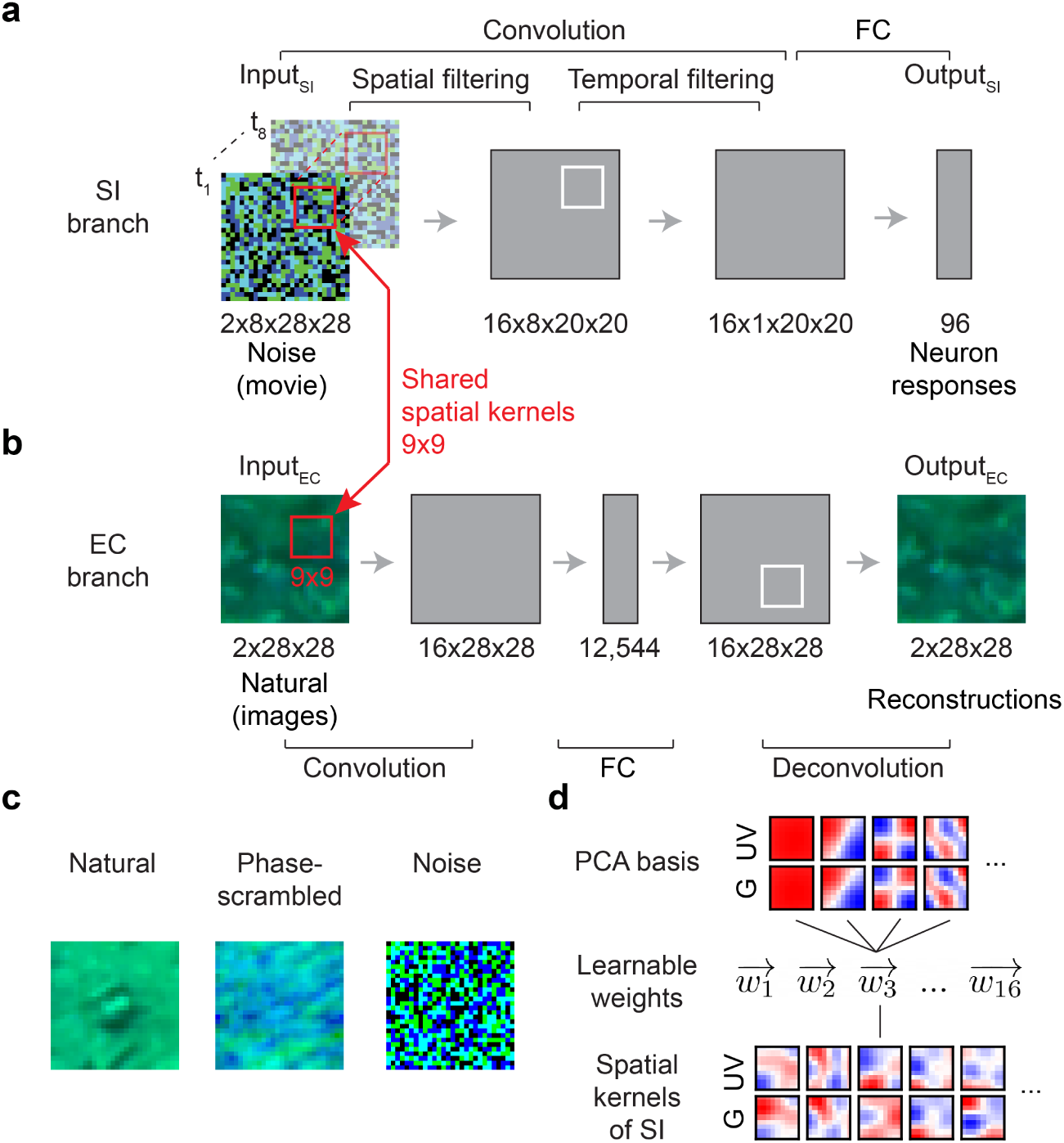
Hybrid model with shared spatial filters. **a,b**. Schemata of SI model (a) and EC model (b) from Qiu et al. (38). The SI model branch consists of spatial and temporal convolutional layers, a fully connected (FC) layer and a nonlinear layer (see Methods). The EC model branch is a convolutional autoencoder, consisting of an encoder and a decoder network. In the hybrid model, the two branches were trained in parallel with shared spatial filters (red). Input_SI_: 8-frame UV-green noise (*t*_1_ … *t*_8_); Output_SI_: predicted GCL cell Ca^2+^ responses; Input_EC_: UV-green natural images; Output_EC_: reconstructed Input_EC_. **c**. Example for the different inputs (natural images, phase-scrambled natural images, and noise) for the EC branch in hybrid models (*hybrid-natural, hybrid-pha-scr, hybrid-noise*). **d**. Using PCA filters as basis vectors for spatial convolutional filters of the SI model; *SI-PCA* learned 16 weight vectors (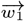 … 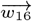) with same vector length as the number of PCA basis elements.

The *EC branch* was trained to efficiently reconstruct input stimuli (i.e., natural scenes) from a constrained latent representation. For this branch, we used a convolutional autoencoder network that we published before (for details, see (38) and Methods). Also in the EC branch, we enforced smooth filters by using L2 regularization, and limited the bandwidth by adding Gaussian noise and imposing L1 regularization on the hidden activations. The latter regularization also encourages sparse representations.

In the *hybrid model*, we implemented interactions between the two branches by shared filters (symbolized by red circle in Fig. 1b). Both branches were trained in parallel, with a weighted sum of their respective losses (*L*_*SI*_ and *L*_*EC*_) used as optimization objective. By changing the weighting of the two losses, we were able to control the relative contribution of two branches on shaping the shared filters, and test our hypothesis to which degree efficient representations of natural scenes improve neural predictions (Fig. 2a,b). Specifically, weight *w* was used to define the hybrid model’s loss function as *L*_*Hybrid*_ = *w* · *L*_*SI*_ + (1 − *w*) · *L*_*EC*_ (Methods). For *w* = 1, the EC branch had no influence on the shared filters and, hence, the hybrid model behaved like the pure SI model. Conversely, for *w* = 0, the SI branch had no influence on the shared filters and, hence, the hybrid model behaved like the pure EC model. Thus, the smaller the weight, the more the EC branch contributed to shaping the filters.

To evaluate the influence of stimulus statistics on neural response predictions, we fed not only natural stimuli to the EC branch, but also phase-scrambled natural stimuli as well as noise. We refer to these models as *hybrid-natural, hybrid-pha-scr* and *hybrid-noise* (Fig. 2c). Moreover, to examine whether the performance improvements could be attributed to simple low-pass filtering, we trained SI networks using spatial convolutional filters composed from different numbers of basis functions derived from principle component analysis (PCA) on natural images (Fig. 2d), or the discrete cosine transform (DCT). These models are referred to as *SI-PCA* and *SI-DCT* networks.

To train the SI branch of our hybrid framework, we recorded somatic Ca^2+^ responses from populations of cells in the ganglion cell layer (GCL) of the *ex-vivo* mouse retina to 9-minute long noise stimuli using two-photon imaging (Fig. 3a; Methods; (43, 44)). The GCL contains the RGCs, which represent the retina’s output neurons and form in the mouse about 40 parallel feature channels to higher visual brain areas (reviewed in (23)). RGCs gain their specific response properties by integrating upstream input from distinct sets of bipolar cells and amacrine cells. Note that the GCL also contains some “displaced” amacrine cells (dACs; (43, 45)). If not indicated otherwise, we did not distinguish between these two GCL cell classes in our datasets. The noise stimulus contained two chromatic components (UV, green) matching the spectral sensitivities of mouse photoreceptors (46). We used the data of n=96 GCL cells that passed our quality criteria (Methods) to fit a pure SI model with factorized spatial and temporal convolutional filters, whose predictive performance served as our baseline (Fig. 3b left).

**Fig. 3.**
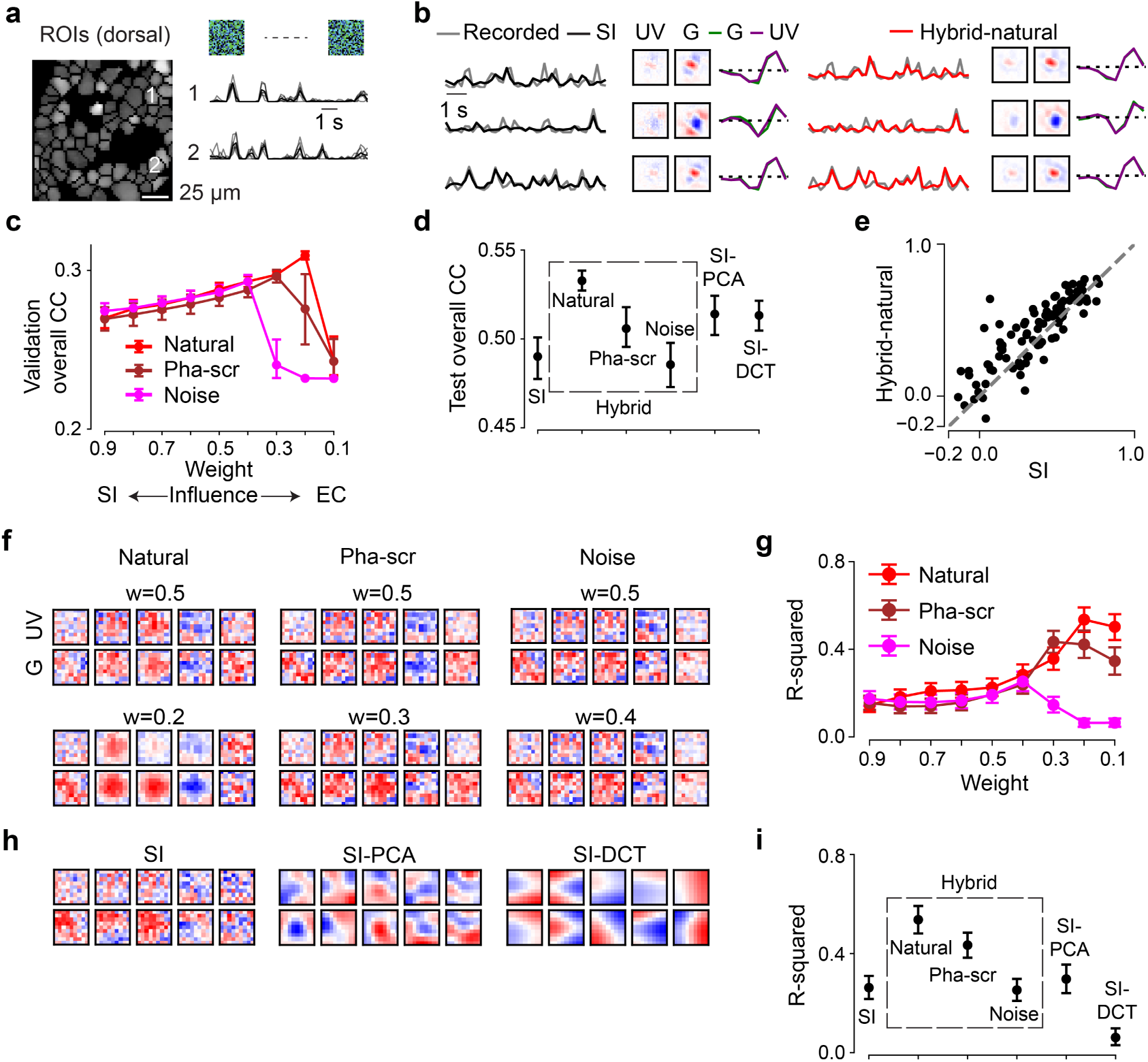
Neural encoding tasks benefit from natural scene statistics. **a**. Region-of-interest (ROI) mask of one recording field in dorsal retina (left) and mean Ca^2+^ responses (black) of exemplary ROIs in response to 6 repeats of noise stimuli (single trials in gray). **b**. Three representative GCL cell responses (gray) to the noise stimulus (cf. Fig. 2a, left), together with predictions of best performing models on test data (black, SI; red, hybrid w/ natural scenes as input to the EC path, i.e., Input_EC_), and learned spatiotemporal receptive fields (RFs) visualized by SVD. **c**. Model performance (linear correlation coefficient, CC; mean for n=10 random seeds per model) based on validation data for hybrid model with natural scenes (red), with phase-scrambled scenes (brown), or with noise (magenta) as Input_EC_, and for different weights. **d**. Best performance (mean for n=10 random seeds per model) based on test data for SI, SI-PCA (16 bases), SI-DCT (4 bases), hybrid-natural (*w*=0.2), hybrid-pha-scr (*w*=0.3) and hybrid-noise (*w*=0.4; p<0.0001 for SI vs. hybrid-natural, p=0.0085 for SI-PCA vs. hybrid-natural, p=0.0011 for hybrid-natural vs. hybrid-pha-scr, two-sided permutation test, n=10,000 repeats). **e**. Scatter plot for model predictions based on test data for hybrid-natural (*w*=0.2) vs. SI at one random seed, with each dot representing one neuron. **f**. Representative spatial filters (shared convolutional filters) for hybrid models with different Input_EC_ and different weights. Upper: with *w*=0.5; lower: with optimal *w* (see (c)) for hybrid models. **g**. Mean R-squared of fitting a 2D Gaussian to spatial filters (cf. (f)), for hybrid model with natural scenes (red), with phase-scrambled scenes (brown), or with noise (magenta) as Input_EC_, and for different *w* (n=10 random seeds per model). **h**. Representative spatial filters (shared convolutional filters) for SI, SI with PCA filters (16 bases) and SI with DCT filters (4 bases). **i**. Mean R-squared of fitting a 2D Gaussian to the spatial filters for one chromatic stimulus channel (green; n=10 random seeds per model; p<0.0001 for SI vs. hybrid-natural, p<0.0001 for SI-PCA vs. hybrid-natural, p=0.0074 for hybrid-natural vs. hybrid-pha-scr, two-sided permutation test, n=10,000 repeats). Error bars in (c),(d),(g),(i) represent 2.5 and 97.5 percentiles obtained from bootstrapping.

### Neural system identification benefits from natural scene statistics

First, we measured the predictive performance of the hybrid-natural model on the validation data (for hyperparameter tuning) by systematically varying the relative impact of the two branches by changing the weight *w*. We found that the performance steadily increased with increasing EC influence (i.e., decreasing *w*) up to an optimum (peaking at *w* = 0.2; Fig. 3c, red), after which the SI had too little influence on the shared filters and the performance dropped. Note that the correlation values for the validation data are relatively low because these predictions were calculated on a single-trial basis (Methods).

Next, we replaced the natural input to the EC pathway by phase-scrambled scenes (*hybrid-pha-scr*) and white noise across space and chromatic channels (*hybrid-noise*). Like for the hybrid-natural model, the performance of the two control models also increased with increasing EC influence up to a certain point, peaking at *w* = 0.3 and *w* = 0.4 for hybrid-pha-scr and hybrid-noise, respectively (Fig. 3c). This indicates that when incorporating EC, all hybrid model versions showed some improvement up to certain *w* values, before performance sharply declined.

To test to what extent simple low-pass filtering contributes to the performance improvement observed for the hybrid-natural model, we quantified the performance of two additional SI models, one with PCA and the other one with DCT bases. By varying the number of bases used, we found a maximum in predictive performance at 16 and 4 bases for SI-PCA and SI-DCT (zig-zag ordering), respectively (Suppl. Fig. S1b).

Finally, to compare the performance on the test data across models, we picked for each model, the *w* or number of bases with the best predictive performance for the validation data. We found that the hybrid model with natural inputs to the EC branch attained the best performance among all tested models (Fig. 3d,e). The hybrid-natural model’s superior performance compared to the hybridpha-scr model suggests that the benefit of learning natural scene statistics extends beyond second-order statistics such as the 1*/f* power spectrum of natural images. Nevertheless, the hybrid-pha-scr model performed better than the hybrid-noise version, pointing at a general benefit of learning second-order statistics in the EC branch. Moreover, the hybrid-natural model was consistently better than low-pass filtering control models (*SI-PCA* and *SI-DCT*), suggesting that simple low-pass filtering does not fully explain the benefits of sharing kernels with the EC branch trained to efficiently represent natural stimuli. Together, our results suggest that normative network regularization — in particular, based on natural statistics — can improve the performance of neural SI models.

### Hybrid models with natural inputs learn the most biologically-plausible filters

To confirm that our hybrid models capture the properties of the recorded cells, we estimated their RFs (Fig. 3b; Suppl. Fig. S1f; Methods). Indeed, we found that the models learned antagonistic center-surround RFs with biphasic temporal kernels, reminiscent of RGC RFs found in other studies (2, 43). To get insights to which degree our models resembled biological vision systems, we next investigated the internal representations by analyzing the filters of the models’ subunits (18, 47). To this end, we compared the shared spatial convolutional filters between our tested models. As neurons in the retina and further upstream in the early visual system often feature smooth, Gaussian or DoG shaped RFs (e.g., (43, 48, 49)), we considered models with such shared filters as more biological plausible than those with other filter organizations.

Interestingly, while the learned neuronal RFs were quite consistent between models (cf. Fig. 3b), their shared spatial filters differed considerably (Fig. 3f,h). When using natural images in the EC branch (*hybrid-natural*), filters indeed became smoother and more Gaussian-shaped, which may be a result of the regularization by the EC branch on the SI branch and which may have contributed to the performance improvement of predicting responses. This effect persisted though reduced when phase-scrambled images were used (*hybrid-pha-scr*). Moreover, for smaller *w* values (i.e., stronger EC influence), Gaussian-shaped filters became more frequent in the hybrid-natural but not in the hybrid-noise model (Fig. 3f, upper vs. lower row). For the SI models with PCA or DCT basis, we found all filters to be smooth as they profited from low-pass filtering of the respective transformation. However, compared to the hybrid-natural model, their filters were less frequently Gaussian-shaped (Fig. 3h).

To quantify these findings, we fitted 2D Gaussian functions to the filters and measured the goodness of the fit via the coefficient of determination (R-squared; Methods). Notably, for all three hybrid models, the *w* with the best Gaussian fit was the same *w* that also resulted in the best response predictive performance (*w* = 0.2, *w* = 0.3, and *w* = 0.4 for *hybrid-natural, hybrid-pha-scr*, and *hybrid-noise*, respectively; Fig. 3g). The filters of the hybrid-natural model resembled smooth 2D Gaussians more than for any other model (Fig. 3i), including SI-PCA and SI-DCT. The difference of fit quality between hybrid-natural vs. hybrid-pha-scr and hybrid-pha-scr vs. hybrid-noise may be related to higher-order statistics and second-order statistics of natural scenes, respectively.

Taken together, our comparisons of the hidden spatial representations suggest that natural scene statistics promote latent feature representations akin to transformations in the early visual system.

### Efficient coding increases the data efficiency of system identification

Next, we asked if the observed performance increase in the hybrid-natural vs. the baseline SI model was sensitive to the amount of training data, both with respect to their response predictions (Fig. 4a) and their learned spatial filters (Fig. 4b). To this end, we trained the SI and the hybrid-natural model (*w* = 0.2) with different amounts of data, ranging from 30% to 100%.

**Fig. 4.**
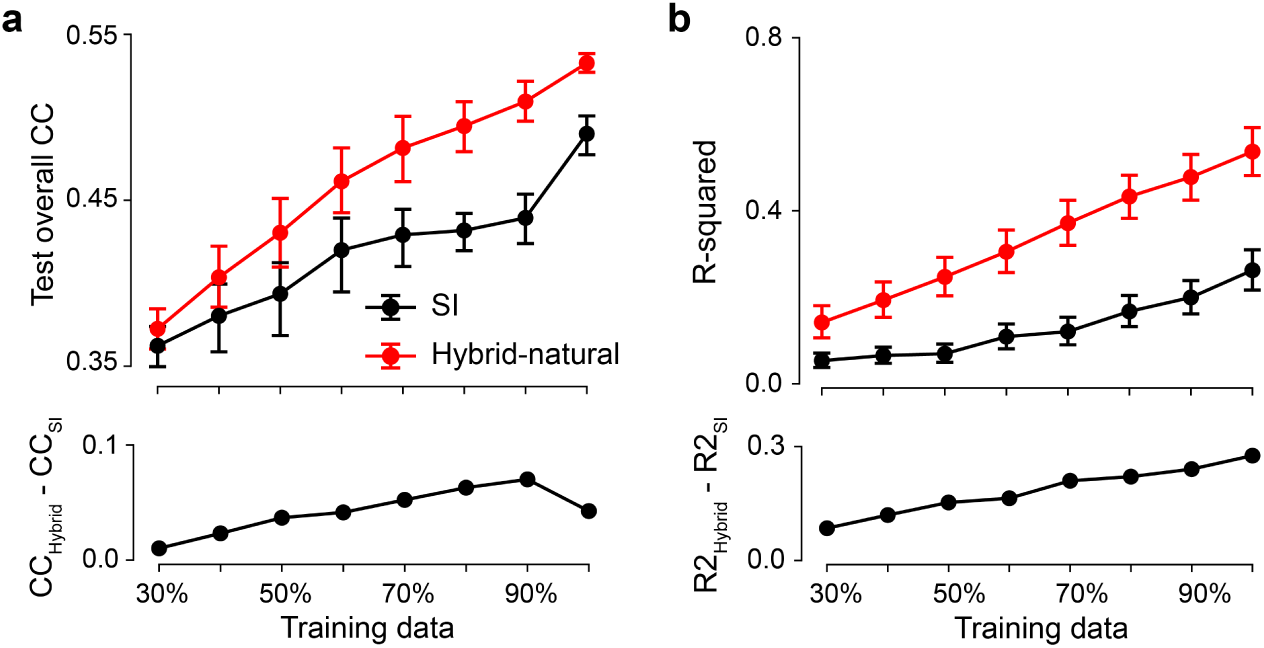
Hybrid-natural models with better data efficiency for neural prediction. **a**. Mean model performance (top) based on test data for SI and hybrid-natural (*w*=0.2; n=10 random seeds) with different training data sizes and mean difference between SI and hybrid-natural (bottom). **b**. Mean R-squared (top) of fitting a 2D Gaussian to spatial filters for green stimulus channel for SI and hybrid-natural (*w*=0.2; n=10 random seeds) with different training data sizes, and the mean difference between R-squared for SI and hybrid-natural (bottom). Error bars represent 2.5 and 97.5 percentiles with bootstrapping.

Not unexpectedly, when more training data was used, predictive performance increased for both models (Fig. 4a top). However, we also found that the performance of the hybrid-natural model was consistently higher than that of the SI model, with the difference becoming significant for ≥60% and peaking at around 90% training data (Fig. 4a bottom). Additionally, for both models the spatial filters became increasingly more Gaussian-like with more data (Fig. 4b). We also observed that the performance difference dropped for large dataset sizes — which, we expect, may be asymptotically near zero in the regime of infinite data.

Together, these results suggest that a hybrid-natural model, which has access to natural statistics, requires significantly less training data than the baseline SI model.

### Hybrid models for testing temporal coding strategies

It has been suggested that early stages of visual processing, rather than encoding a past stimulus, aim at predicting future stimuli in their temporal stream of inputs (24, 50–52). Such a future prediction strategy is thought to require a smaller dynamic range to be encoded than that needed for representing past stimuli (past encoding), and thus allows for lower energy consumption (53, 54). Therefore, we next tested if the neural encoding task would profit even more from natural statistics when spatio-temporal (i.e., 3D) filters were shared between the hybrid model’s two branches. We implemented both strategies — past encoding and future prediction — in the EC branch, and compared their influence on the SI task (55).

We modified the 2D SI model to use spatio-temporal (instead of factorized spatial and temporal) convolutional filters to predict neural responses for 8-frame noise movies (*3D SI model*; Suppl. Fig. S2a). Likewise, we employed spatio-temporal convolutional filters for the EC branch. As before, the two branches of the resulting hybrid model were trained in parallel, but now sharing spatio-temporal filters. In the past encoding case, the EC branch was trained to reconstruct the 7^th^ frame (at *t* − 1) of a continuous 8-frame natural movie clip based on frames at *t* − 7 to *t* (*hybrid-natural-past*; Suppl. Fig. S2b,c). In the future prediction case, the EC branch was trained to predict the 8^th^ unseen frame based on the first 7 frames (*t* − 7 to *t* − 1) of the clip (*hybrid-natural-future*; Suppl. Fig. S2d left).

Like for the 2D models, we varied *w* or the number of bases and then selected the best model for each condition (*3D SI, hybrid-natural-past, hybrid-natural-future*, and *3D SI-PCA*) based on validation performance. We next quantitatively compared the different models using the test data (Fig. 5a,b; Suppl. Fig. S3c). We found that the 3D SI-PCA model outperformed the 3D SI model, presumably because the former profited from the low-pass filtering of the PCA transformation. Importantly, both hybrid models displayed a better performance than the 3D SI-PCA model. While the hybrid-natural-past model performed slightly better than its hybrid-natural-future counterpart, this difference was not statistically significant. In summary, both the past encoding and future prediction strategy in the EC branch turned out to be equally beneficial for the neural encoding task and, as before, the benefit extended beyond low-pass filtering effects. However, no performance increase was achieved with respect to the 2D hybrid-natural model (Fig. 5b vs. Fig. 3d).

**Fig. 5.**
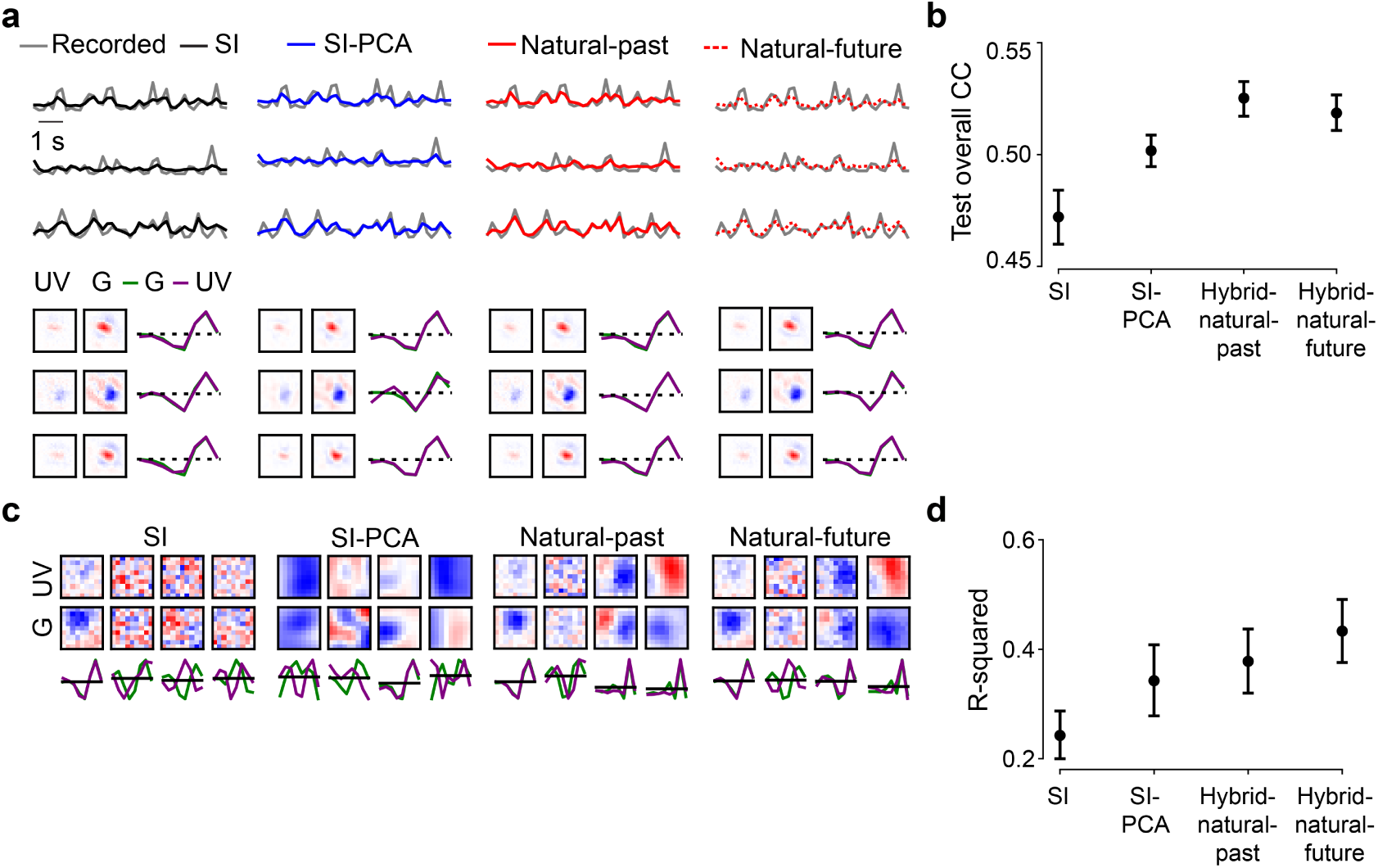
Past encoding or future prediction strategies using 3D shared filters perform equally well. **a**. Top row: Responses of three exemplary GCL cells to 5-Hz noise stimulus (gray) and predictions of best performing models on test data (black, SI; blue, SI with PCA filters; red solid, hybrid for encoding the past; red dotted, hybrid for predicting the future). Bottom row: Respective learned RFs of the three cells (visualized by SVD). **b**. Mean model performance based on test data for SI, SI-PCA (128 bases), hybrid-natural-past, and hybrid-natural-future (both *w*=0.4; n=10 random seeds; p<0.0001 for SI vs. hybrid-natural-past, p=0.0005 for SI-PCA vs. hybrid-natural-past, p=0.2563 for hybrid-natural-past vs. hybrid-natural-future, two-sided permutation test, n=10,000 repeats). **c**. Representative shared spatial and temporal filters of 3D models (n=1 random seed, visualized by SVD; temporal kernels for UV and green stimulus channels indicated by purple and green, respectively). **d**. Mean R-squared of fitting a 2D Gaussian to shared spatial filters (for green stimulus channel; n=10 random seeds per model; p=0.0003 for SI vs. hybrid-natural-past, p=0.4356 for SI-PCA vs. hybrid-natural-past, p=0.1895 for hybrid-natural-past vs. hybrid-natural-future, two-sided permutation test, n=10,000 repeats). Error bars in (b),(d) represent 2.5 and 97.5 percentiles with bootstrapping.

We also analyzed the shared spatio-temporal filters using the same metric as for the 2D case, which assesses the similarity between spatial filters (after performing a low-rank decomposition of 3D shared filters into spatial and temporal components; see Methods) and smooth 2D Gaussians (Fig. 5c,d). Again, we found higher R-squared values for the hybrid models and the 3D SI-PCA model compared to the baseline SI case. Note that here, the 3D SI-PCA model did not significantly differ from the two hybrid models, possibly due to a large number of bases (*n* = 128 vs. *n* = 16 in the 2D case).

Next, we asked if the fact that we did not see a significant advantage of 3D over 2D could be due to the relatively slow (5 Hz) noise stimulus, which may drive insufficiently temporal properties of the GCL cell responses. Therefore, we recorded a new dataset (*n* = 64 cells) in which we presented a 30-Hz dense noise stimulus and used it with the 3D hybrid models. Like for 5-Hz noise, hybrid-natural-past and hybrid-natural-future models performed similarly on the validation data, with a peak in performance at around *w* = 0.7 (Suppl. Fig. S4a), as well as on the test data, where they were significantly better than the 3D SI model (Suppl. Fig. S4b). Moreover, both 3D hybrid models learned shared filters with similar R-squared values, which were significantly higher than that of the 3D SI model (Suppl. Fig. S4c). But again, the 3D models performed only equally well compared to the 2D models.

In summary, the hybrid-natural models achieved a higher performance for different noise stimuli (5-Hz vs. 30-Hz) and different shared filter organizations (2D vs. 3D) than all other tested models. Therefore, it is likely that their superior predictive performance for neuronal responses and their more biologically plausible filters resulted from the EC branch having access to natural statistics.

### Direction-selective neurons benefit more than others from hybrid models

The retina encodes the visual scene in a number of features that are represented by the more than 40 different types of RGC whose outputs are relayed in parallel to higher visual centers in the brain (43, 56–59). Thus, we next asked, if access to natural statistics allows our hybrid models to predict some cell types better than others (Fig. 6). Earlier, it has been shown that motion-relevant properties emerge in the efficient coding framework for both past encoding and future prediction approaches (55). Therefore, we employed our 3D hybrid models (cf. Fig. 5) and focused on direction-selective (DS) cells (43, 60).

**Fig. 6.**
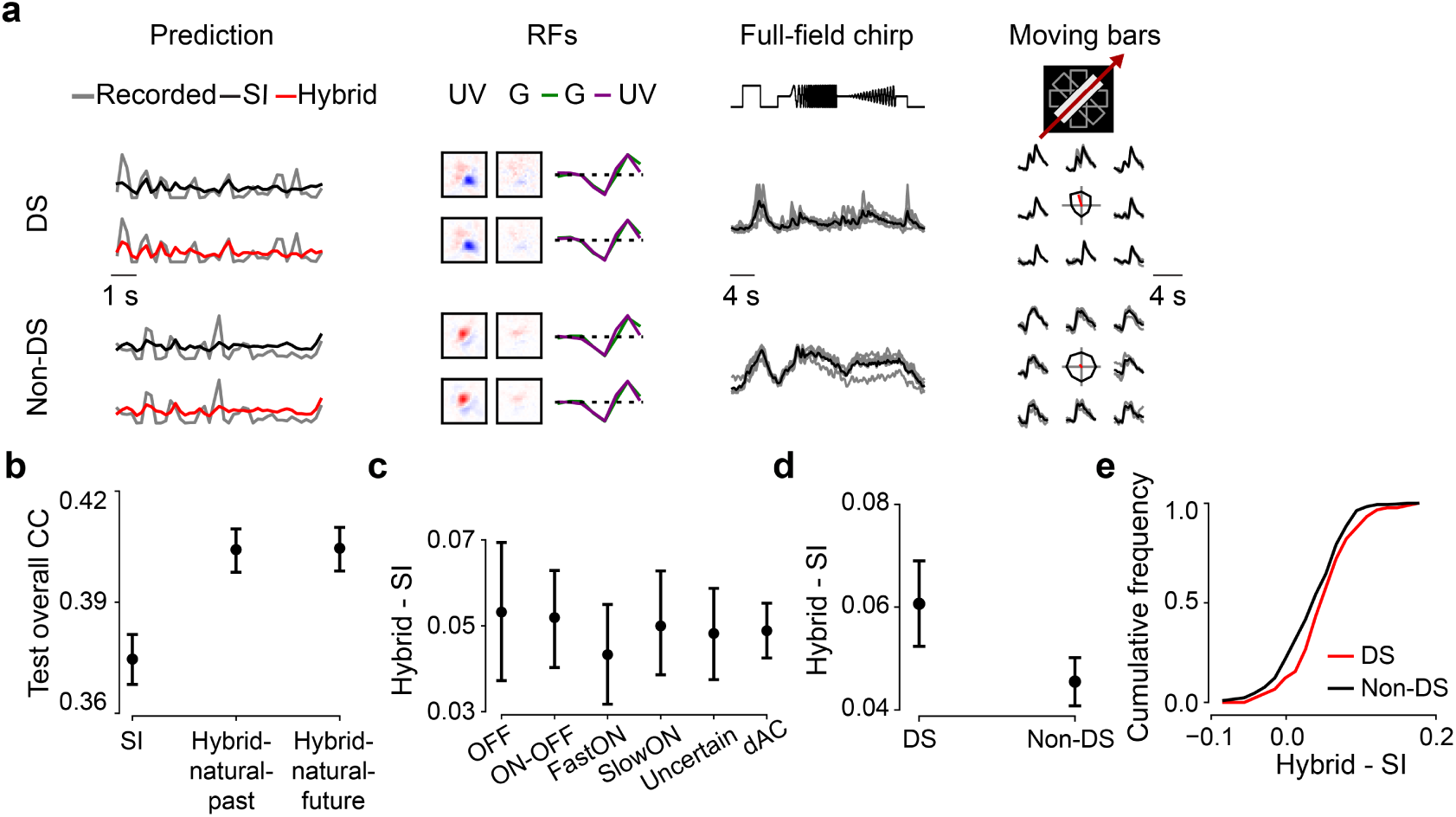
Direction-selective (DS) neurons benefit more from hybrid models. **a**. Recorded (gray) and predicted (black, SI; red, hybrid-natural-past; response amplitude scaled with a constant 1.5 for better visualization) responses to noise, RFs, as well as full-field chirp responses and moving bar responses (gray, single trials; black, means) of representative DS and non-DS cells. Note that the RFs were dominated by UV stimulus channel because cells were recorded in ventral retina (see Methods). **b**. Mean model performance based on test data for SI, hybrid-natural-past and hybrid-natural-future (both *w* = 0.7; n=10 random seeds per model; trained with responses of n=427 GCL cells to 5-Hz noise stimulus; p<0.0001 for SI vs. hybrid-natural-past, p=0.9307 for hybrid-natural-past vs. hybrid-natural-future; two-sided permutation test, n=10,000 repeats). **c**. Difference in mean performance between hybrid-natural-past and SI based on test data for 6 broad functional groups of GCL cells (35 OFF, 59 ON-OFF, 49 fast-ON, 38 slow-ON, and 64 uncertain RGCs, as well as 145 dACs; see Methods and Results; n=10 random seeds per model). **d**. Like (b) but for n=90 DS and n=300 non-DS cells. **e**. Cumulative histogram of difference in mean prediction between hybrid-natural-past (*w* = 0.7) and SI on test data for DS (red) and non-DS cells (black), at one particular seed. Error bars in (b)–(d) represent 2.5 and 97.5 percentiles with bootstrapping.

For this analysis, we used a set of n=427 GCL neurons, whose responses were recorded not only to the 5-Hz noise stimulus (for training the models) but also to full-field chirp and moving bar stimuli. The latter two stimuli (Fig. 6a) enabled us to identify the functional type of each recorded GCL neuron (43) using a cell type classifier (see Methods; Suppl. Fig. S5).

To explore cell type-specific effects, we chose a dataset size (30% of total recording time) for which the synergy between neural SI and EC was particularly pronounced. As expected, we found that both hybrid networks (*hybrid-natural-past* and *hybrid-natural-future*) performed significantly better than the SI model, with no significant difference between the two hybrid models (cf. Fig. 5b, Suppl. Fig. S4b).

First, we evaluated if any of the broader functional groups of GCL cells profited more from natural statistics than others. For this, we sorted the cells into 6 groups based on their response polarity (ON vs. OFF) and transience, and based on whether they were RGCs or dACs (for group sizes, see Fig. 6 legend). For all 6 groups, the hybrid models showed a better predictive performance than the SI model (Fig. 6b). However, no significant differences were observed between any pair of groups (p>0.05 for all pairwise comparisons, two-sided permutation test, n=10,000 repeats; Fig. 6c) and the two hybrid models (p>0.05 for all pair-wise comparisons; Suppl. Fig. S6a).

Next, we grouped the cells into DS (p<0.05, direction tuning using a permutation test; n=90) and non-DS cells (n=300) based on their moving bar responses (Fig. 6a right). Note that n=37 neurons were excluded as they did not pass the quality check for chirp and moving-bar responses (Methods). We found that the predictive performance for DS cells was significantly higher than that of the non-DS cells for both hybrid-natural-past (Fig. 6d,e; p=0.0027) and hybrid-natural-future (Suppl. Fig. S6b,c; p=0.0042). To test whether this performance difference was merely due to different signal-to-noise ratios in DS vs. non-DS cells, we compared their response quality indices (*QI*; Methods). While DS cells had significantly higher *QI* values for moving-bar responses (*QI*_*bar*_) than non-DS cells, we did not find any significant difference between the two groups with respect to their noise (*QI*_*noise*_) or chirp responses (*QI*_*chirp*_; Suppl. Fig. S6e-g). These results suggest that DS cells benefit more from the EC branch of the hybrid models than non-DS cells, partially consistent with earlier findings ((55); see also Discussion).

In summary, efficient coding of natural statistics served as a beneficial normative regularization for all types of mouse GCL cells and in particular DS cells, suggesting the potential role of motion statistics in the natural environment on shaping neuronal response properties.

## Discussion

In this study, we asked if access to natural scene statistics can help predicting neural responses. To address this question, we combined system identification (SI, (3)) and efficient encoding (EC, (25)) methods into a normatively regularized (hybrid) modeling framework. Specifically, we used models that efficiently represent natural scenes recorded in the mouse’ habitat to regularize models that predict retinal responses to visual stimuli. We analyzed such hybrid models with shared spatial filters, and found that natural images as input to the EC branch indeed improved the performance in predicting retinal responses and allowed the model to generate filters that resembled RFs found in the early visual system. These improvements extend beyond those gained by simple low-pass filtering or using second-order statistics of the natural scenes. Our hybrid models with shared spatio-temporal filters performed similarly well as those with shared spatial filters, independently of whether they used a past encoding or a future prediction strategy. Notably, predictions for DS cells in the mouse retina improved the most in the hybrid models with natural input. In summary, our results suggest that sourcing information about an animal’s environment — e.g., through hybrid SI-EC models — helps building more predictive and biologically-plausible models of neuronal networks. More generally, our findings lend support to the idea that knowledge of natural statistics is already encoded in sensory circuits.

### Hybrid models improve data efficiency

The difference in predictive performance between the hybrid and the baseline SI model was significant and it depended on the amount of available data, indicating that our hybrid modeling approach increased data efficiency. We note that both the stimulus (dense noise) and the neural model system (retinal neurons) present much easier SI problems than, for instance, predicting more nonlinear neural responses to natural stimuli (18, 61). For those more challenging problems at downstream visual areas, where neural response functions and, hence, the neural prediction tasks, become more complex (62), the data efficiency of a hybrid approach and the improvement from natural scene statistics may be even higher.

### Biological plausibility and temporal coding principles in hybrid models

The biological plausibility of most learned models was positively correlated with their predictive performance except some indeterminacy for SI-DCT models, suggesting that more biologically plausible filters increased performance. Note that we used the filters’ similarity to smooth 2D Gaussian functions as a measure of biological plausibility, following the assumption that RFs in the retina (and at early downstream stages of the visual system) often feature smooth, Gaussian-like structure (43, 48, 49). However, a deep, systematic understanding of artificial and neuronal networks and their hidden representations likely calls for other methods besides of filter inspection (discussed in (63)).

As the natural environment is not static, we also created hybrid models that acknowledge the time domain by sharing spatio-temporal filters. Surprisingly, both variants — past encoding and future prediction — behaved quite similar. However, in the stand-alone EC models (that is only the respective EC branch), the temporal components of the filters learned by the future prediction were much more diverse than those of past encoding (Suppl. Fig. S2c,d right). Interestingly, the differences between temporal filter of these stand-alone EC models decreased with the incorporation of the neural prediction task in the hybrid models.

The filter diversity in our 3D hybrid models is reminiscent of earlier findings by Chalk and colleagues (2018), who reported the emergence of filters sensitive to motion direction and motion speed in their past encoding and future prediction EC models, respectively. However, in contrast to their results, we did not see a difference between our hybrid-past and hybrid-future models with respect to motion-sensitive filters: Both of them performed better in predicting responses of DS vs. non-DS cells. Further work is needed to understand that partial (mis)match between our work and that by Chalk et al., and why specifically DS cells profited from both our 3D hybrid models.

### Hybrid models of retinal signal processing

It has been suggested that natural stimuli drive more diverse neural responses, and more complex feature transformations are required to determine the respective stimulus-response functions ((18, 64), but also see (65)). Therefore, one future direction may be to record retinal activity while presenting natural movies (e.g., from (38)) and use it as input for the SI branch of the hybrid model. Finding a more pronounced performance improvement compared to the baseline SI model would support the notion that the noise stimulus we used in this study may have indeed limited the benefits from the EC branch (see above). Neural data to natural stimuli would also allow us to revisit our hybrid models with respect to the prediction of motion sensitive cells and the differences between our results and those from earlier work ((55); see above). Furthermore, such data may be useful for characterizing model generalization (domain transfer, see e.g., (61, 64)) by using responses to natural stimuli as unseen test data with a hybrid model trained with cell responses to noise stimuli.

For our current analysis, we used broad group assignments (e.g., FastON RGCs), which include several functional types of RGC (e.g., ON-step, ON-transient, ON-high-frequency etc; (43)) or dACs, but did not detect any differences in performance gain except for the DS neurons. Still, it is possible that distinct types of RGC profit more than others from the EC branch of our hybrid models. For example, the so-called W3 RGCs, for which the best stimulus found so far is a small dark moving spot (66), may not be “designed” to efficiently represent natural stimuli but rather to extract survival-relevant features (i.e., detecting aerial predators). Here, we could build models with different normative regularization or tasks (i.e., detecting predators in images of the sky) and would expect that this RGC type profits little from efficiently encoding natural statistics in the hybrid model. Studying coding strategies across RGC types could contribute an important biological perspective to the perennial debate between efficient coding (67) and feature detection (56) proponents.

### Normative network regularization as a framework for studying neural coding

In this study, we regularized the filters of a SI model with a normative EC model to predict visually-evoked responses of cells in the retina. Some forms of such normative regularization have also been discussed and/or applied in earlier work. For example, Deneve and Chalk (68) discussed the relations between SI (encoding) models and EC, and argued that the latter may promote shifting the focus in SI from the single-cell to the population level. The integration of stimulus-oriented approaches (such as EC) for discriminative tasks (such as object recognition) was proposed by Turner et al. (15). Later, Teti et al. (69) employed sparse coding with lateral inhibition in simulations of neuronal activation in visual cortex. More recently, Młynarski et al. (41) presented a probabilistic framework combining normative priors with statistical inference and demonstrated the usefulness of this approach for the analysis of diverse neuroscientific datasets. However, their work was rather conceptual, with the datasets they used being either simulated or low-dimensional. Notably, they tested their framework on pre-fit retinal RFs, but not directly on actual RGC stimulus-response data. Compared to their framework, our method does not require marginalization across all parameter space to estimate optimality and could be applied to more general or complex inference problems. Hence, our work not only provides further evidence to the feasibility of combining coding principles for identification of neural response properties on high-dimensional data, it also demonstrates the benefits of leveraging natural scene statistics for neural prediction. However, compared to the framework by Młynarski et al., with our approach it is more difficult to conduct rigorous statistical tests of normative theory.

We expect that our hybrid modeling strategy may also work for different processing stages along the early visual pathway (and potentially other modalities, e.g., sound). This said, however, one needs to keep in mind that different stages along the visual pathway have different tasks and constraints, and, thus, likely incorporate different efficient coding principles: For instance, the retinal hardware is space-limited and has to encode visual features in view of a bottleneck with limited bandwidth (optic nerve), whereas the primary visual cortex has comparably abundant resources which might serve for accurate probability estimation for behavioral tasks, such as novelty detection (discussed in (24, 70)). It is also worth to note that different visual processing stages (such as primary visual cortex vs. higher visual areas, or adaptation of visual coding to different behavioral states) may benefit from the hybrid modeling to a different degree, as efficient coding approaches learn filters that may be more relevant to stimulus-related features, but not high-level behavior goals (see discussion in (15)). Additionally, it would be interesting to compare our hybrid models with SI models regularized with other behavioral tasks such as object recognition (e.g., (11)) or predator detection (see above) for neural predictions along the ventral visual stream.

There is a long tradition of using SI models (reviewed in (3)) in predicting the responses of neurons to a great variety of stimuli (e.g., (2, 4, 18, 19, 71, 72)). Our results demonstrate how the EC hypothesis can be successfully leveraged as normative regularization for the identification of neural response properties. More generally, using EC as a flexible tool to impose regularization on modeling, the hybrid framework offers an opportunity to test different coding principles and unsupervised learning objectives with regards to experimental data for understanding neuronal processing.

## Materials and Methods

### Animal procedures and retinal activity recordings

#### Animal procedures

All animal procedures were performed in accordance with the law governing animal protection issued by the German Federal Government (Tierschutzgesetz), approved by the governmental review board (Regierungspräsidium Tübingen, Baden-Württemberg, Konrad-Adenauer-Str. 20, 72072 Tübingen, Germany). We used n=5, 5-9 weeks old female C57BL/6 mice (wild-type; JAX 000664, Jackson Laboratory, USA). Due to the exploratory nature of our study, we did not use any statistical methods to predetermine sample size, nor did we perform blinding or randomization.

Animals were housed under a standard light-dark (12h:12h) cycle. All procedures were carried out under very dim red illumination (>650 nm). Prior to the start of the experiment, animals were dark-adapted for ≥ 1 h, then anesthetized with isoflurane (Baxter, Germany), and killed by cervical dislocation.

The eyes were enucleated and hemisected in carboxygenated (95% O_2_, 5% CO_2_) artificial cerebrospinal fluid (ACSF) solution containing (in mM): 125 NaCl, 2.5 KCl, 2 CaCl_2_, 1 MgCl_2_, 1.25 NaH_2_PO_4_, 26 NaHCO_3_, 20 glucose, and 0.5 l-glutamine (pH 7.4). Next, the retina was flat-mounted onto an Anodisc (#13, 0.2 *µm* pore size, GE Healthcare, Germany) with the ganglion cell layer (GCL) facing up. To uniformly label the GCL cells, bulk electroporation was performed with the fluorescent

Ca^2+^ indicator Oregon-Green BAPTA-1 (OGB-1; Invitrogen, Germany), as described earlier (44, 73), using 4-mm plate electrodes (CUY700P4E/L, Xceltis, Germany) and 9 pulses (∼ 9.2 V, 100 ms pulse width at 1 Hz). After electroporation, the tissue was immediately moved to the microscope’s recording chamber, where it was continuously perfused with carboxygenated ACSF at 36^°^C and left to recover for ∼ 30 min before recordings started. Additionally, Sulforhodamine-101 (SR101, Invitrogen, Germany) was added to the ACSF (∼ 0.1 *µ*M final concentration) to visualize blood vessels and identify damaged cells.

#### Two-photon Ca^2+^ recordings and light stimulation

We recorded light stimulus-evoked Ca^2+^ signals in GCL cells of the explanted mouse retina using a MOM-type twophoton (2P) microscope (74, 75) from Sutter Instruments (purchased from Science Products, Germany), as described earlier (43, 44). In brief, the microscope was powered by a mode-locked Ti: Sapphire laser (MaiTai-HP DeepSee, Newport Spectra-Physics, Germany) at 927 nm. Two detection pathways allowed simultaneously recording of OGB-1 and SR101 fluorescence (HQ 510/84 and HQ 630/60, respectively; both Chroma/AHF, Germany) through a 16x water immersion objective (CFI75 LWD×16 /0.8W, DIC N2, Nikon, Germany). A custom-written software (ScanM, by M. Müller and T.E.) running under IGOR Pro 6.3 for Windows (Wavemetrics, USA) was used to acquire time-lapsed (64×64 pixels) image scans at a frame rate of 7.8125 Hz. Higher resolution images were acquired using 512×512 pixel scans. Additionally, to register the scan field positions, the outline of the retina and the optic disc were traced.

The retinas were presented with color noise stimulus using a visual stimulator tuned to the spectral sensitivities of mice (76). This stimulus consisted of independent binary dense noise (28×28 pixel frames, each pixel covering (0.83^°^)^2^ of visual angle) in the UV and green stimulator channels at 5 or 30 Hz. The stimulus contained 5 different training sequences (96 s each) interspersed with 6 repeats of a 10 s test sequence (Suppl. Fig. S1a).

In total, we used three data sets for modeling: (*i*) responses of n=96 GCL neurons to 5-Hz noise recorded in dorsal retina (n=2 eyes); (*ii*) responses of n=427 GCL neurons to 5-Hz noise recorded ventrally (n=5 eyes); in this dataset, we also presented two other stimuli: a full-field chirp (700 *µ*m in diameter) and a moving bar stimulus (300×1,000 *µ*m bright bar moving at 8 directions at 1 mm/s). The responses to these latter stimuli were used to functionally classify the recorded GCL neurons (43). (*iii*) n=64 GCL neurons to 30-Hz noise recorded ventrally (n=2 eyes). Note that all cell numbers are after quality control (see below).

#### Data preprocessing and analysis

For each cell, we calculated a quality index (*QI*, with 0 ≤ *QI* ≤ 1) for its responses to each stimulus type as follows:

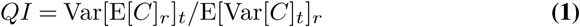

where C is a t-by-r response matrix (time samples, *t*, by repetitions, *r*). The higher *QI*, the more reliable the response and the higher the signal-to-noise ratio. For the noise stimulus, *QI*_*noise*_ was determined based on the test sequence responses. For the following analysis, we only used cells with *QI*_*noise*_ *>* 0.25; in case chirp and moving bar responses were also recorded, neurons had to fulfill *QI*_*chirp*_ *>* 0.35 or *QI*_*bar*_ *>* 0.6 to be included.

In case of the noise stimulus, we preprocessed each cell’s Ca^2+^ signal by Z-normalizing the raw traces and matching sampling frequency of the recording (7.8125 Hz) to the stimulus frequency (5 or 30 Hz) via linear interpolation. Then, the traces were detrended using a high-pass filter (*>* 0.1 Hz) and their 1^st^ order derivatives were calculated, with negative values set to zero. We used the average of a cell’s responses to the 6 test sequence repeats as ground truth. Excluding the test sequences, we had per cell a total of 480 s of data, of which we used 440 s (∼ 91%) for training and the remaining 40 s (∼ 9%) for validation (i.e., to pick the hyperparameters of the SI model, see below).

For chirp and moving bar responses, we first detrended the traces and then normalized them to [0, 1] (44). Using these responses, the cells were classified to different functional groups (43) using RGC type classifier (see below).

To estimate the directional tuning from the moving bar responses, we first performed singular value decomposition (SVD) on the mean response matrix, resulting in a temporal and a directional component. We then summed the directional vectors in 2D planes and used the resulting vector length as direction selectivity index. Next, by shuffling trial labels and computing the tuning curve for 1,000 times (permutation test), we got the null distribution (no directional tuning). The percentile of true vector length was used as p-value of directional tuning (43). Here, we considered cells with *p <* 0.05 as direction-selective (DS) and the remaining ones as non-DS.

#### RGC type classifier

To predict the functional type of GCL cells, we used a Random Forest Classifier (RFC; (77)), which was trained on a published mouse dataset (43). In that study, features were extracted from the responses to different visual stimuli (e.g., chirp and moving bar) and used to cluster GCL cells into 32 RGC types and 14 additional dAC types. Here, we learned a mapping *f* from response features (20 features from responses to chirp, *ϕ*_*chirp*_ and 8 features from responses to moving bar stimulus, *ϕ*_*mb*_) and two additional parameters Θ = {*θ*_*soma*_, *θ*_*DS*_} to functional cell type labels *L* by training a RFC for the dataset from (43):

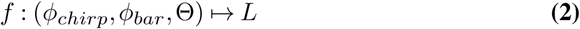

where *θ*_*soma*_ denotes soma size to distinguish between alpha and non-alpha RGC types and *θ*_*DS*_ denotes p-value of permutation test for direction selectivity to distinguish between DS and non-DS RGC types.

We fit the RFC on a subset of data from (43) and validated its performance on a held-out test dataset. The classifier had a prediction accuracy of ∼ 76% on a held-out test dataset (Suppl. Fig. S5). To apply the trained classifier to our newly recorded dataset, we projected the RGC responses (normalized to [− 1, 1]) into the feature space described in (43) by computing the dot product between the response and the feature matrices. We used the RFC implementation provided by the python package `scikit-learn` (78) to train the classifier.

### 2D models

#### Stand-alone SI model (2D)

As baseline model to predict the responses of neurons to the noise stimulus, we employed a stand-alone SI model (supervised learning), in which we used factorized spatial and temporal convolutional filters (Fig. 2a; (79, 80)). This SI model consisted of one spatial convolutional layer (16×2×1×9×9, output channels x input channels x depth x image width x image height), one temporal convolutional layer (16×16×8×1×1, with 8 stimulus frames preceding an event), and — after flattening the spatial dimension — one fully connected layer (FC; 96×6,400, output x input channels), followed by an exponential function. No padding was used. The loss function was defined as:

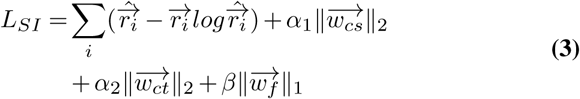

Here, the first term is the Poisson loss between predicted responses 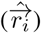 and ground truth 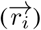 (with *i* denoting the neuron index), the second term is the L2 penalty on the weights of the spatial convolutional filters 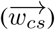 with hyperparameter *α*_1_, the third term is the L2 penalty on the weights of temporal convolutional filters 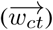 with hyperparameter *α*_2_, and the last term is the L1 penalty on the FC layer 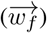 with hyperparameter *β*.

After performing a grid search for the three hyperparameters, we picked *α*_1_ = 10, *α*_2_ = 10, *β* = 1*/*16 which yielded the best performance on the validation data. After training, we estimated the neurons’ spatio-temporal RF filters by computing gradients for each neuron, starting with a blank image sequence as input. These gradients represent the first-order approximation of the input that maximizes the neuron’s activation (6). For visualization, we extracted the spatial and temporal RFs via SVD.

As a metric of biological plausibility, we calculated the coefficient of determination (R-squared; [0, 1]) of fitting 2D Gaussian distributions to the spatial (component of) the convolutional filters. We set the R-squared value to 0 if the sigma of the fitted Gaussian was larger than the size of the filter (i.e., 9 pixels). We calculated this fit quality for the filter of the chromatic channel with the dominant response. Because the mouse retina is divided into a more green-sensitive dorsal and a more UV-sensitive ventral retina (e.g., (44)), this meant that for dorsal neurons we only determined the R-squared for filters for the green stimulus channel, and for ventral neurons for the UV stimulus channel.

#### SI-PCA model (2D)

The spatial convolutional filters of the SI-PCA model were composed from PCA basis functions (*W*). The model was trained to learn the weights of these basis functions. The filters were produced by performing PCA transformation on natural images recorded in mouse habitats (38):

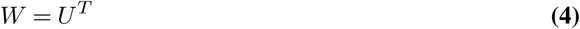

where *U* contains the eigenvectors of the covariance matrix of the centered data in each column.

For example, when using 4 PCA bases, the shape of learnable weight matrix was 16×4 (channel number x basis number), the shape of PCA bases was 4×2×1×9×9 (basis number x chromatic channel x depth x image width x image height), and the resulted spatial filter had the shape of 16×2×1×9×9. We varied the number of used basis (hyperparameter) and selected the one which achieved the best performance on validation data (Suppl. Fig. S1b; Suppl. Fig. S3b).

#### SI-DCT model (2D)

For the SI-DCT model, its spatial convolutional filters were composed from DCT basis functions, which were defined as:

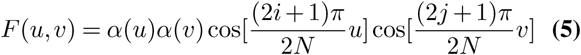

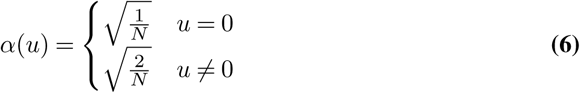

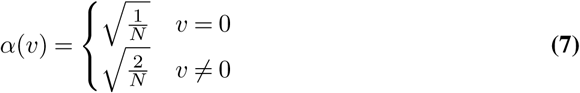

where *i* and *j* denote pixel index of the input image (size (*N, N*)); *u* and *v* denote DCT coefficient index of the DCT filter. Here, we employed DCT basis functions for one-channel gray images and thus used different bases for each chromatic channel. For example, when using 4 DCT bases, the shape of learnable weight matrix was 16×4×2 (channel number x basis number x chromatic channel), the shape of basis function was 4×1×9×9 (basis number x depth x image width x image height), and the resulted spatial filter had the shape of 16×2×1×9×9. Like for SI-PCA, we varied the number of used basis and picked the one which achieved the best performance on validation data (Suppl. Fig. S1b).

#### Stand-alone EC model (2D)

We used a similar EC model architecture (convolutional autoencoder) and loss function as in (38). The model’s encoder contained a single convolutional layer (with weights denoted 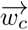) followed by a rectified linear unit (ReLU) function, one FC layer, and another ReLU function. The decoder contained one FC layer, one ReLU function, a single deconvolutional layer (with weights denoted 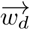), and a hyperbolic tangent (tanh) function to map back to the original data range ([−1, 1]).

As a measure of reconstruction quality, we used mean squared error (MSE; (37, 38)). Gaussian noise was added to the encoder output for redundancy reduction (37, 81, 82) and an L1 penalty (hyperparameter *β*) was imposed to its activation 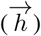 for sparse readouts (37, 81, 83). We also applied L2 regularization on the convolutional and deconvolutional layers to encourage the learning of smooth filters (42, 84, 85). We used 16 9×9 convolutional and deconvolutional filters. The activation tensor (16×28×28, output channel x image width x image height) following the first convolutional layer was flattened to a one-dimensional vector with 12,544 inputs before feeding into the FC layer. The loss function for the EC model was:

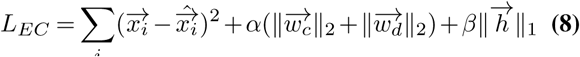

where the first term is the MSE error between the prediction 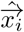 and ground truth 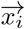 with image index *i*, and the next two terms denote the L2 and L1 penalties.

#### Hybrid model (2D)

The hybrid (semi-supervised) model consisted of a SI and an EC branch (for details on the two models’ architectures, see above). These branches were trained simultaneously, sharing the spatial convolutional filters 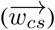. The total loss function of the hybrid model was derived from the loss functions of the two branches as follows:

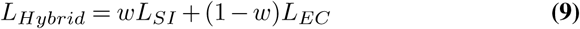

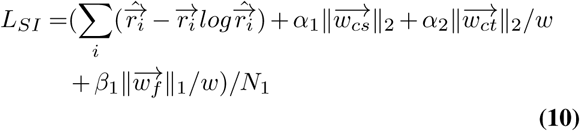

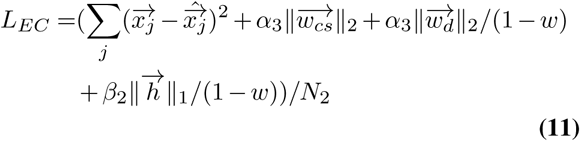

Here, *i* and *j* denote neuron and image index, respectively; 905 *N*_1_ and *N*_2_ the number of neurons and images, respectively. The weight (*w*, with 0 ≤ *w* ≤ 1) controlled the impact of each branch’s loss function on the shared spatial filters. Practically, we used *w* = 10^*−*8^ for *L*_*SI*_ and 909 *w* = (1 − 10^*−*8^) for *L*_*EC*_ when *w* = 0 and *w* = 1, respectively. Note that we added *w* to the denominator of the last two terms to maintain the same regularization for 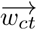 and 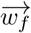 in a stand-alone SI model when varying *w*. For *L*_*EC*_, similar to *L*_*SI*_, we added (1 − *w*) to the denominator of the last two terms to keep the same regularization for 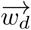 and 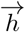 in a stand-alone EC model when varying *w*. We used different data to train the EC branch of the hybrid model: natural images, phase-scrambled natural images and noise. All hybrid models were trained for a maximum of 100 epochs (Suppl. Fig. S1c,d); training was stopped early when the validation loss started decreasing.

Tuning all hyperparameters jointly in a grid search was computationally prohibitive. Hence, for the SI branch, we varied the hyperparameters around those determined for the stand-alone configuration (*α*_1_ = 10, *α*_2_ = 10, *β*_1_ = 1*/*16; see above), while for the EC branch, we varied the hyperparameters systematically around the values (*α*_3_ = 10^3^, *β*_2_ = 1*/*16) used in (38). To tune *w*, we devised a linear search approach by normalizing the loss functions (using *N*_1_ and *N*_2_).

After training the hybrid model, we estimated the spatiotemporal RFs of all neurons using a gradient ascent algorithm (6). We visualized the spatial and temporal component of RFs using SVD (cf. Fig. 3b), and the magnitude of the RF was indicated in the spatial component.

We trained 2D models using all training data (440 s) with a learning rate of *µ* = 10^*−*4^. In case less data were used (i.e., to evaluate data efficiency), we kept all hyperparameters the same as for the full data case but doubled the learning rate. This was done because the stand-alone SI model and the hybrid model could not reach the minimum of validation loss within 100 epochs (when less data were used).

### 3D models

#### Stand-alone SI model (3D)

The 3D SI model consisted of one spatio-temporal convolutional layer (16×2×8×9×9, output channels x input channels x depth x image width x image height; depth varied with the frequency of noise stimuli, n=8 and n=30 for 5-Hz and 30-Hz noise, respectively), and — after flattening all dimension — one FC layer (96×6,400, output channels x input channels; output channel varied with cell numbers n=96, 64 or 427 for different data sets; see above), followed by an exponential function. No padding was used. The loss function was defined as:

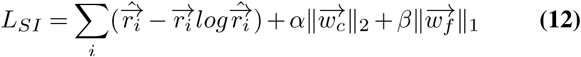

This equation differs from Equation () with respect to the L2 penalty, which is here on the weights of the spatiotemporal convolutional filters 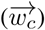 with hyperparameter *α* for the second term. After performing a grid search for the two hyperparameters, we picked *α* = 100, *β* = 1*/*4 which yielded the best performance on the validation data. After training, we estimated and extracted the cells’ spatial and temporal RFs via SVD for visualization.

#### SI-PCA model (3D)

For the 3D SI-PCA models, we applied Equation () to the movie clips (2×8×9×9, chromatic channel x depth x image width x image height; depth varied with the frequency of noise stimuli, n=8 and n=30 for 5-Hz and 30-Hz noise, respectively). Like for 2D SI-PCA models, we varied the number of used bases and picked the number for which the model achieved the best performance on the validation data (Suppl. Fig. S3a).

#### Stand-alone EC model (3D)

The 3D EC models used a sequence of frames from a movie clip as input and featured 3D spatio-temporal convolutional layers (with weights denoted 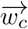) in the encoder. The decoder contained deconvolutional layers with weights 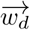. In the past-encoding case, we fed an 8-frame clip (frames at *t* − 7 to *t*) to the model and aimed at reconstructing the 7^th^ frame (at *t* − 1). In the future-prediction case, the goal was to predict the 8^th^ frame (at *t*) with the input being the first 7 frames (*t* − 7 to *t* − 1) of the clip. The loss functions was similar to that given by Equation () except that (*i*) 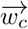 features different a shape (16×2×8×9×9, output channel x chromatic channel x filter depth x filter width x filter height), and (*ii*) *x*_*i*_ denotes the 7^th^ frame for the past encoding and the 8^th^ frame for the future prediction model (Suppl. Fig. S2b,c,d).

#### Hybrid model (3D)

The 3D hybrid models consisted of a SI branch and an EC branch with shared spatio-temporal convolutional filters (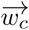; see above). Like for the 2D hybrid models, the total loss function was a weighted sum of losses for the two branches as follows:

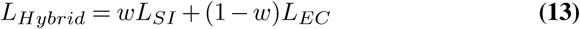

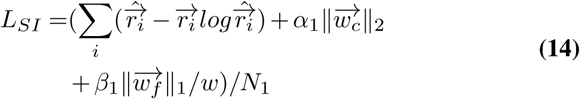

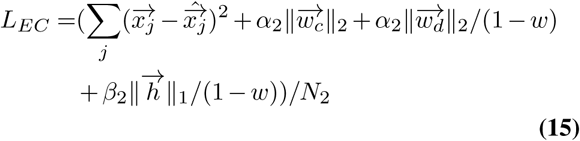

Here, *i* denotes neuron index, *j* movie clip index, *N*_1_ neuron number, and *N*_2_ the number of movie clips. Again, instead of tuning all hyperparameters jointly via a grid search, we varied the hyperparameters around the values determined for the stand-alone SI configuration (*α*_1_ = 100, *β*_1_ = 1*/*4) for the SI branch. For the EC branch, we varied the hyperparameters systematically around the values (*α*_2_ = 10^4^, *β*_2_ = 1*/*16) used in the stand-alone EC models. We then tuned *w* linearly after normalizing the loss functions (using *N*_1_ and *N*_2_). We also visualized the spatial and temporal RF components using SVD (Fig. 5a, bottom).

## Acknowledgments

We thank Matthew Chalk, Dylan Paiton and Katrin Franke for helpful discussions, and Merle Harrer for excellent technical assistance. This work was supported by the German Research Foundation (DFG): SFB 1233, Robust Vision: Inference Principles and Neural Mechanisms, projects 10 and 12, project number: 276693517; and under Germany’s Excellence Strategy EXC 2064/1 (project number 390727645); the European Union’s Horizon 2020 research and innovation programme under the Marie Skłodowska-Curie grant (agreement No 674901); the Max Planck Society (M.FE.A.KYBE0004); and the German Ministry of Education and Research (BMBF; FKZ: 01GQ1002), and the Tübingen AI Center (FKZ: 01IS18039A). The funders had no role in study design, data collection and analysis, decision to publish, or preparation of the manuscript.

## Author Contributions

Conceptualization: Y.Q.; Methodology: Y.Q., D.K., K.S., D.G., L.H., T.S., and T.E.; Data acquisition & curation: K.S.; Formal analysis: Y.Q. with input from D.K., M.B., L.B., and T.E.; Investigation: Y.Q. with input from D.K., K.S., L.B., M.B., and T.E.; Writing – original Draft: Y.Q., D.K., L.B., and T.E.; Writing – review & editing: all authors; Visualization: Y.Q.; D.G. (confusion matrix); Software: Y.Q.; L.H. and D.G. (classifier); Resources: T.S. and T.E.; Supervision: M.B., L.B., and T.E.; Funding acquisition: L.B., M.B., and T.E.

## Data and Code Availability

Data and code would be available upon publication.

---

**Supplemental Fig. S 1.**
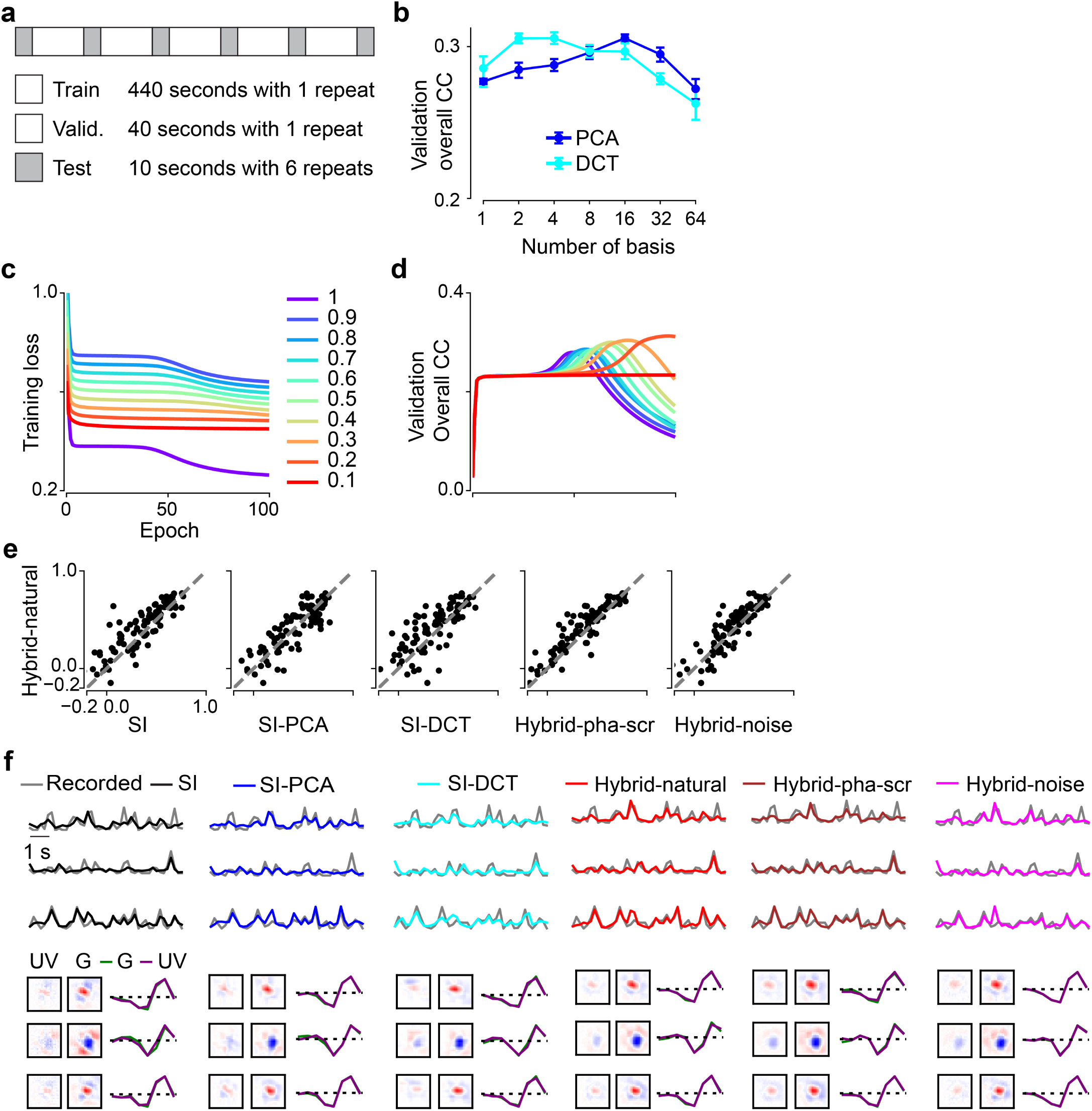
Training of 2D models. **a**. The noise stimulus (9 minutes in total) containing training and validation data (1 repeat) and test data (6 repeats). **b**. Model performance (mean) based on validation data for SI-PCA and SI-DCT with different numbers of basis. SI-PCA and SI-DCT yielded best performance when using 16 and 4 bases, respectively (each model for n=10 random seeds; error bars represent 2.5 and 97.5 percentiles with bootstrapping). **c**. Training loss as a function of training epochs for the hybrid model (Input_EC_, natural scenes) with different weights (*w*), indicated by color (right). **d**. Model performance based on validation data (with linear correlation coefficient as metric) during the hybrid-natural model training with different weights (colors as in (c)). As weight decreased from 1 to 0.2, more training epochs were needed to reach the best performance. The hybrid model performed best for *w* = 0.2. Note that the hybrid model showed a slower change in correlation coefficient (CC) around the peak at *w* = 0.2 (compared to *w* = 1), demonstrating the regularization effects of the EC branch on the hybrid model. **e**. Scatter plots for model predictions based on test data at a particular seed (each dot representing one neuron). Hybrid with natural scenes as input_EC_ (*w* = 0.2) vs. SI, SI with PCA basis (16 bases), SI with DCT basis (4 bases), hybrid-pha-scr (*w* = 0.3) and hybrid-noise (*w* = 0.4). **f**. Upper: Three representative GCL cell responses (gray traces) to noise stimulus together with predictions of the best performing models on test data (black, SI; blue, SI with PCA basis; cyan, SI with DCT basis; red, hybrid w/ natural scenes as input in EC path; brown, hybrid w/ phase-scrambled scenes as input in EC path; magenta, hybrid w/ noise as input in EC path). Lower: Learned spatio-temporal RFs of the example cells, visualized by SVD. Same random seed as in (e).

**Supplemental Fig. S 2.**
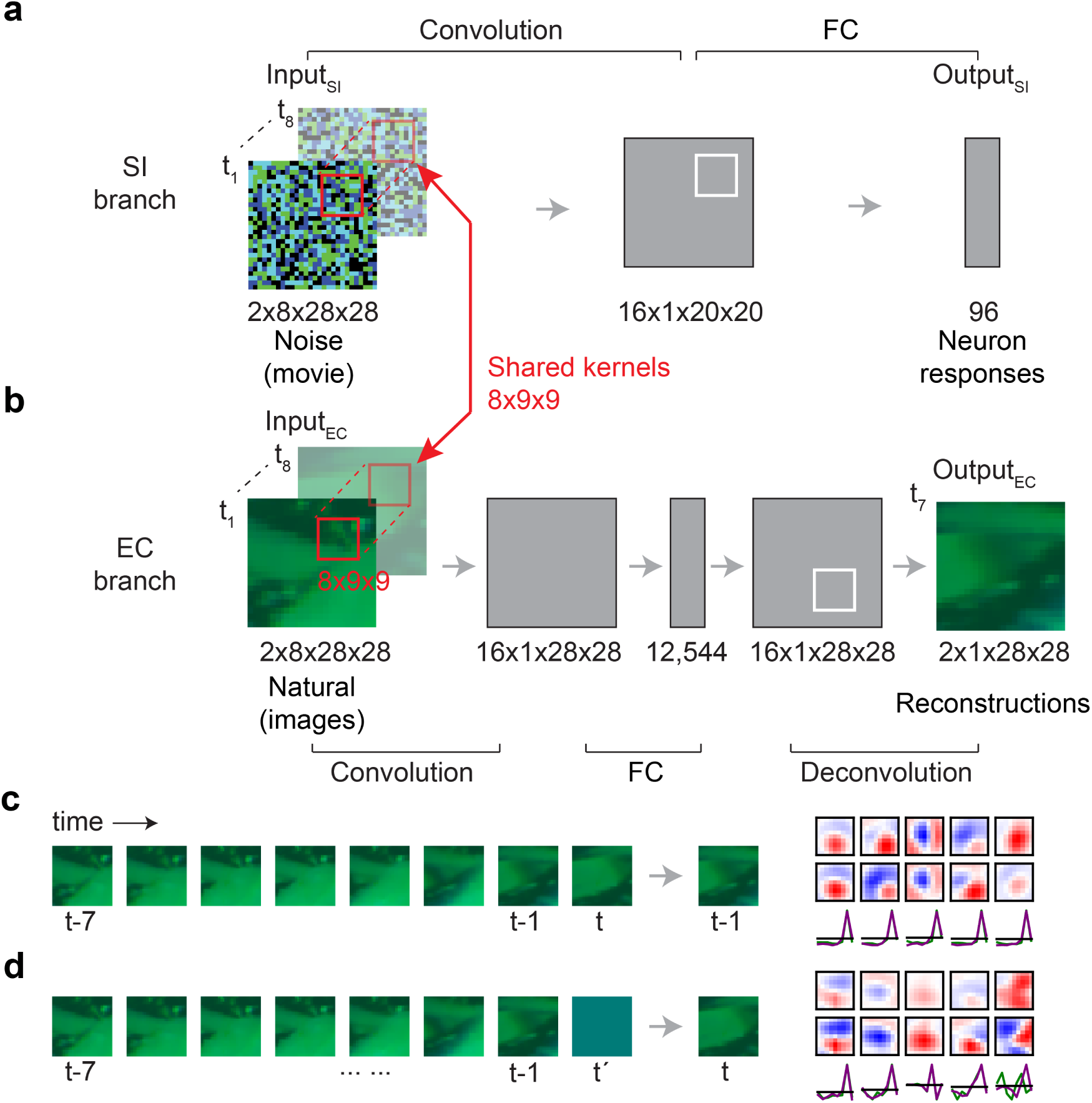
Three-dimensional hybrid networks embedding natural movies. **a,b**. Illustration of SI network (a) with 3D spatio-temporal convolutional filter, and EC network (b), reconstructing the 7^th^ frame (at *t −* 1) based on 8 continuous frames (*t −*7 to *t*; encoding the past, c). Combined as a hybrid network, the two branches were trained in parallel with shared 3D filters (Input_EC_, 8-frame UV-green movie clip; Output_EC_, reconstruction of the 7^th^ frame of Input_EC_). **c**. Example for input/output of the EC model for encoding the past (left; also see b) and exemplary spatio-temporal convolutional filters when using natural movies as input to train the EC model alone (right). **d**. Example for input/output of the EC model for predicting the future, i.e., predicting the 8^th^ frame from the first 7 frames (*t −* 7 to *t −*1) of the clip, and exemplary spatio-temporal filters when using natural movies as input to train the EC model alone. During preprocessing, the 8^th^ frame of input was set to the mean of the first 7 frames, for UV and green channel, respectively. Note that for stand-alone EC models, all temporal components of filters for past encoding were very similar while those for future prediction were much more diverse.

**Supplemental Fig. S 3.**
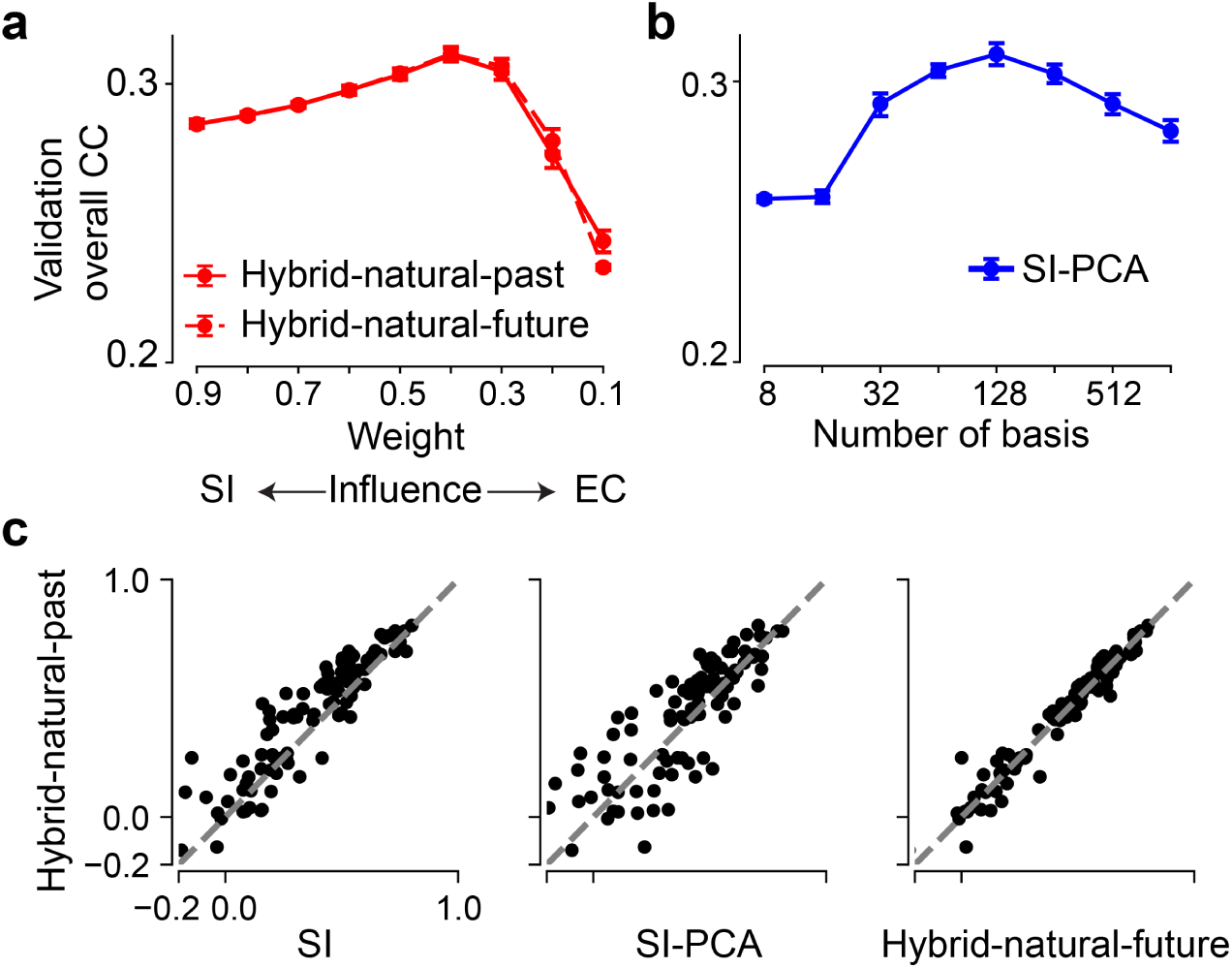
Training of 3D hybrid models. **a,b**. Model performance (mean) based on validation data for hybrid models w/ natural movies as input_EC_ (a), applying past encoding (hybrid-natural-past) or future prediction (hybrid-natural-future) and for different weights, and for the SI-PCA model (b) with different numbers of basis (each model for n=10 random seeds). **c**. Scatter plots for model predictions based on test data at a particular seed (each dot representing one neuron). hybrid-natural-past (*w* = 0.4) vs. SI, SI-PCA (128 PCA bases) and hybrid-natural-future (*w* = 0.4). Error bars in (a)–(b) represent 2.5 and 97.5 percentiles with bootstrapping. Both 3D hybrid models performed similarly, with a peak in predictive performance on the validation data at around *w* = 0.4 (a). This value of *w* was higher than for the 2D hybrid models (*w* = 0.2; cf. Fig. 3c). We also examined the low-pass filtering effects on the 3D SI model by using PCA filters (3D SI-PCA) and varying the number of basis (b). Like for the 2D case when varying the number of basis, we found a maximum in performance on the validation data at 128 bases, which was larger than the 16 bases in the 2D case (cf. Suppl. Fig. S1b).

**Supplemental Fig. S 4.**
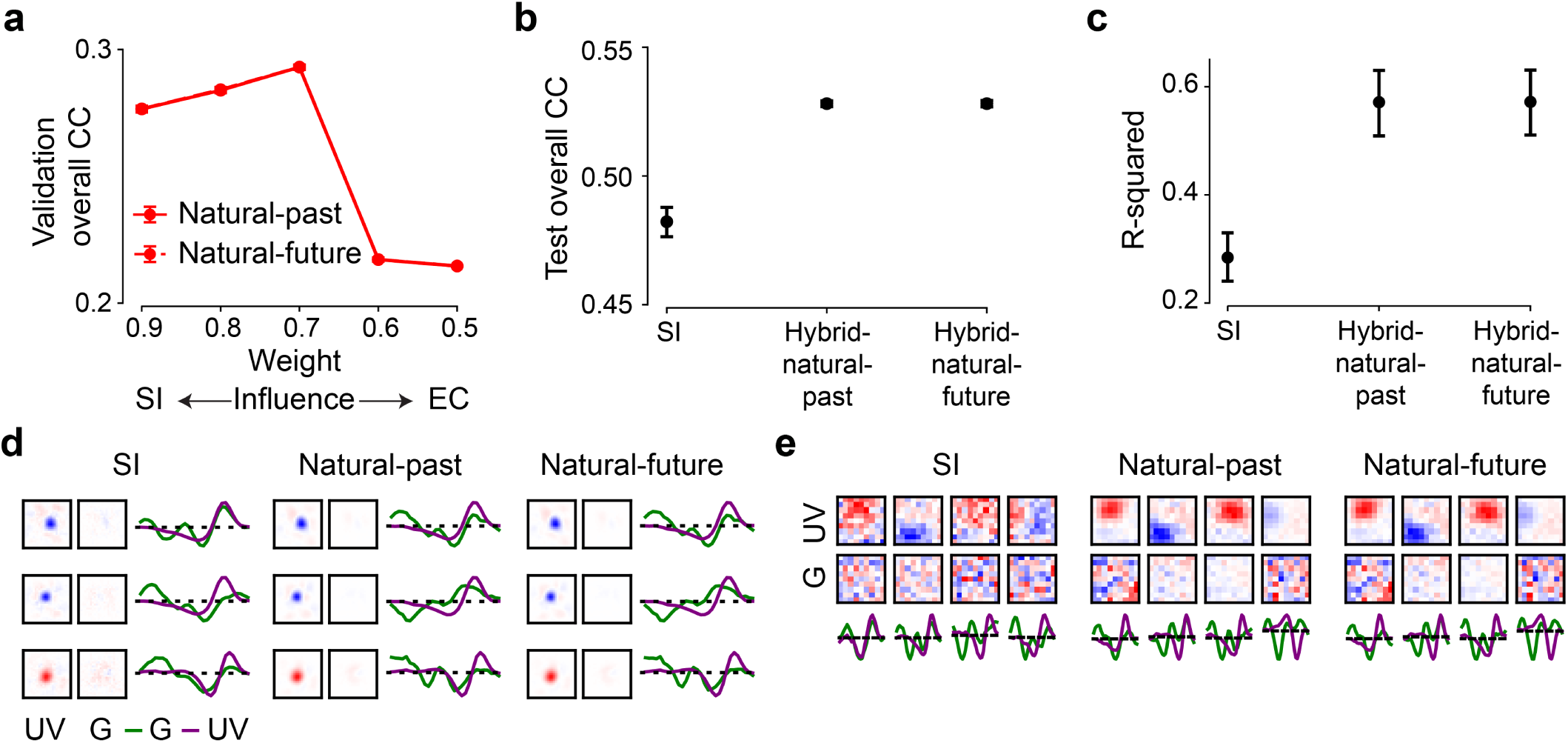
Hybrid model for encoding neuronal responses to 30-Hz dense noise. To test hybrid models for different stimuli, we recorded neuronal responses to the 30-Hz dense noise in the ventral retina. We yielded n=64 neurons after quality control (Methods), which were used to train the SI and hybrid networks. **a**. Model performance (mean) based on validation data for hybrid models (w/ natural movies as input_EC_), applying encoding-past (hybrid-natural-past) or predicting-future (hybrid-natural-future) and for different weights. Each model for n=10 random seeds. Both models with similar performance for all weights, peaking at *w* = 0.7. **b**. Model performance (mean) based on test data for SI, hybrid-natural-past (*w* = 0.7) and hybrid-natural-future (*w* = 0.7). Each model for n=10 random seeds. The two hybrid models had better performance with smaller standard deviation compared the SI model (p<0.0001 for SI and hybrid-natural-past, p=0.9992 for hybrid-natural-past and hybrid-natural-future; two-sided permutation test, n=10,000 repeats). **c**. R-squared (mean) of fitting a 2D Gaussian to all the spatial filters in UV stimulus channel (each model for n=10 random seeds; p<0.0001 for SI and hybrid-natural-past, p=0.9888 for hybrid-natural-past and hybrid-natural-future; two-sided permutation test, n=10,000 repeats). **d**. Learned spatio-temporal filters of the three representative cells, visualized by SVD. Note that because all neurons in this data set were recorded in the ventral retina, their responses were dominated by the UV channel. Different temporal filters in the UV channel were observed for these neurons (cf. the very similar temporal filters in the green channel for neurons’ responses to 5-Hz noise in Fig. 3b, Fig. 5a lower). **e**. Exemplary shared spatial and temporal filters of 3D models, visualized by SVD and for one random seed. Temporal: UV and green channels indicated by purple and green lines, respectively. Error bars in (a)–(c) represent 2.5 and 97.5 percentiles with bootstrapping.

**Supplemental Fig. S 5.**
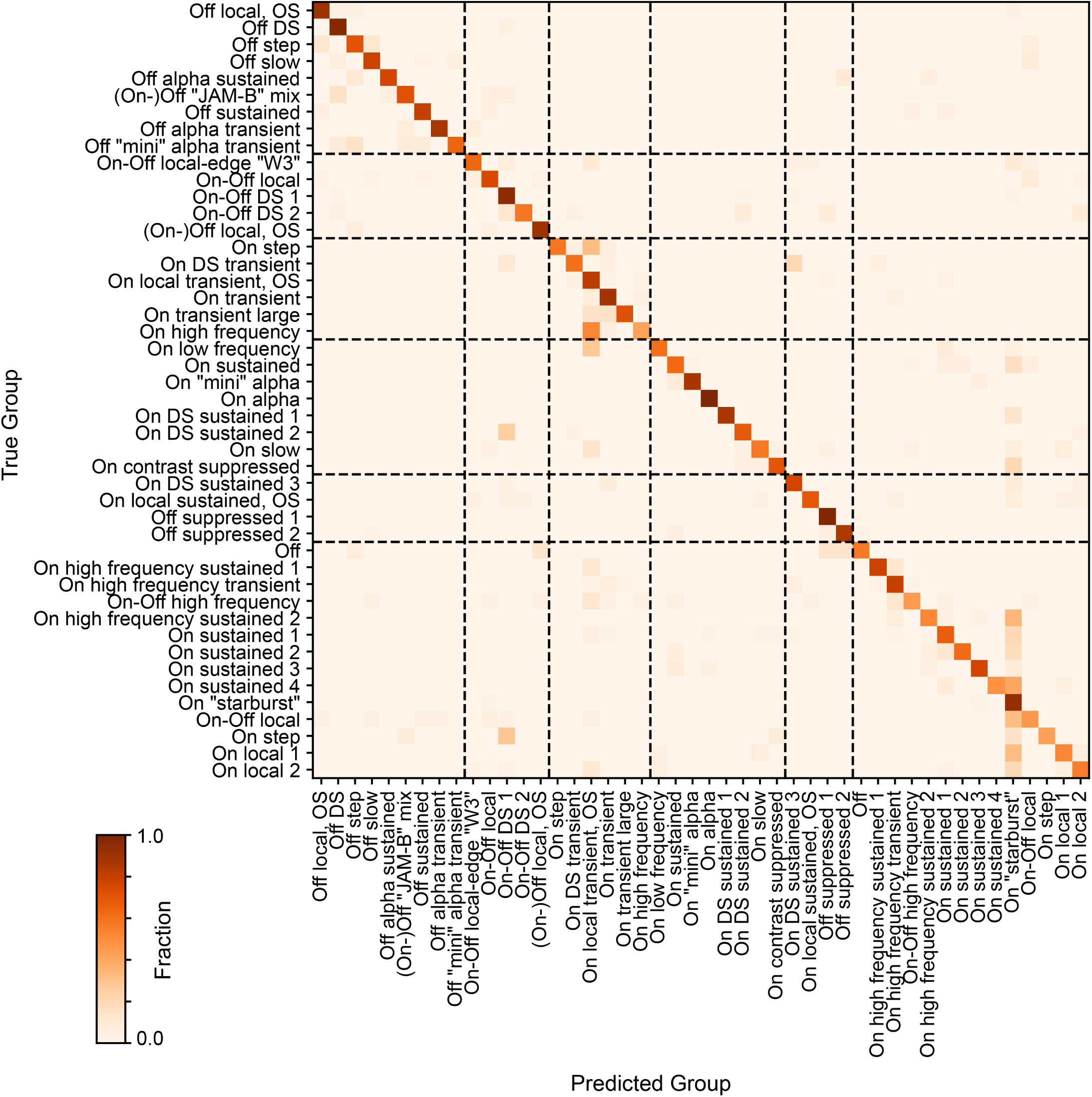
Confusion matrix for a trained random forest classifier. Normalized confusion matrix (true cell types against predicted cell types) for a trained random forest classifier evaluated on a test dataset (for details, see Methods). Dotted line indicates separation of 6 broad functional cell groups (43).

**Supplemental Fig. S 6.**
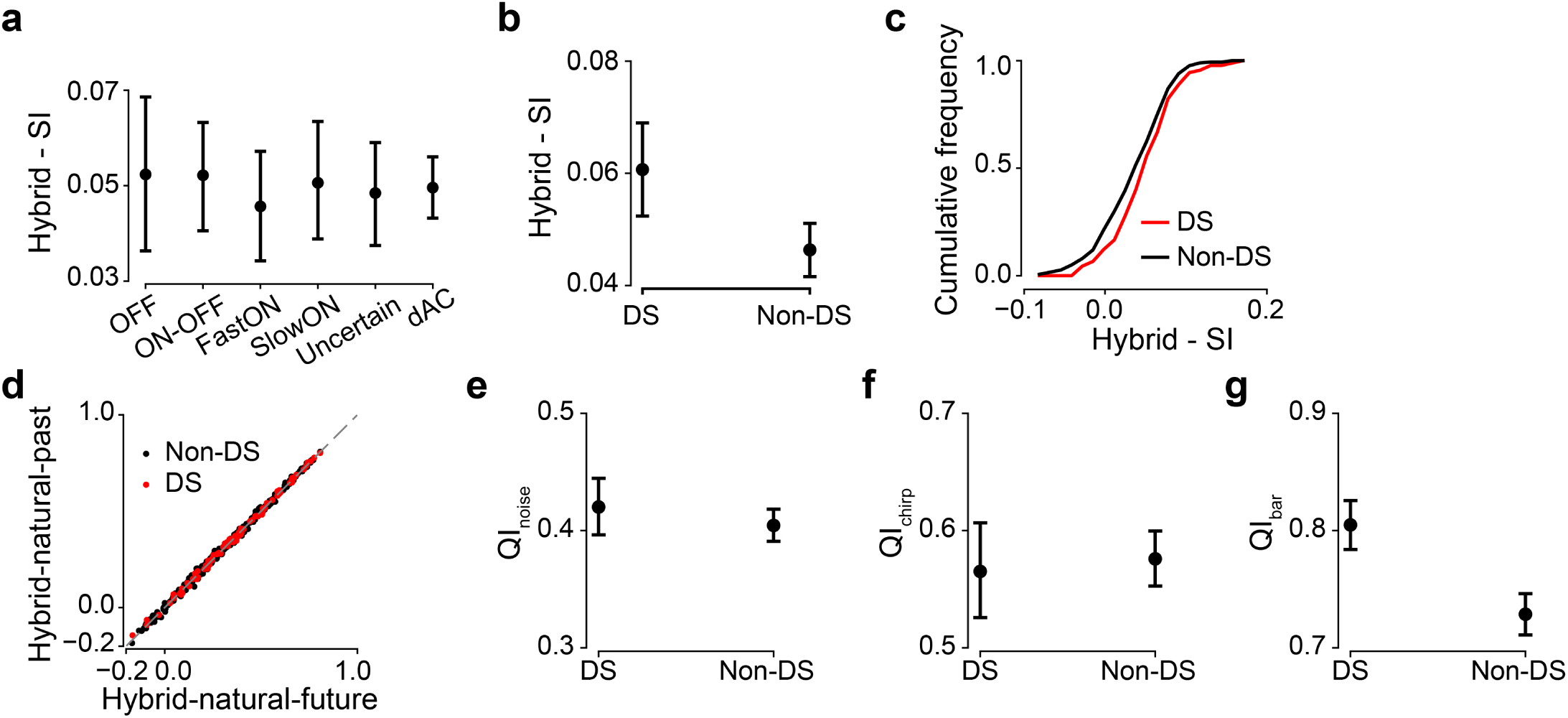
Hybrid model for different cell types. **a**. Performance difference (mean) between hybrid-natural-future and SI based on test data for different cell types (each model for n=10 random seeds). **b**. Performance difference (mean) between hybrid-natural-future and SI based on test data for DS and non-DS cells (each model for n=10 random seeds). **c**. Cumulative histogram of model prediction difference between hybrid-natural-future (*w* = 0.7) and SI on test data, for DS (red) and non-DS cells, at one particular seed. **d**. Scatter plots for model predictions based on test data at a particular seed (each dot representing one neuron) for DS and non-DS cells and hybrid-natural-past (*w* = 0.7) vs. hybrid-natural-future (*w* = 0.7). Note that the predictions of two hybrid models were similar for most of neurons. **e**. Quality index (mean) for DS and non-DS cells based on responses to the repeated test sequences in the noise stimuli (p=0.2881, two-sided permutation test, n=10,000 repeats; for details, see Methods). **f**. Like (e) but for chirp responses (p=0.6714, two-sided permutation test, n=10,000 repeats). **g**. Like (e) but for bar stimulus responses (p<0.0001, two-sided permutation test, n=10,000 repeats). Error bars in (a),(b),(e)-(g) represent 2.5 and 97.5 percentiles with bootstrapping.

## Bibliography

1. Ian H Stevenson and Konrad P Kording. How advances in neural recording affect data analysis. Nature neuroscience, 14(2):139–142, 2011.

2. EJ Chichilnisky. A simple white noise analysis of neuronal light responses. Network: Computation in Neural Systems, 12(2):199–213, 2001.

3. Michael C-K Wu, Stephen V David, and Jack L Gallant. Complete functional characterization of sensory neurons by system identification. Annu. Rev. Neurosci., 29:477–505, 2006.

4. Jonathan W Pillow, Jonathon Shlens, Liam Paninski, Alexander Sher, Alan M Litke, EJ Chichilnisky, and Eero P Simoncelli. Spatio-temporal correlations and visual signalling in a complete neuronal population. Nature, 454(7207):995–999, 2008.

5. Vasilis Marmarelis. Analysis of physiological systems: The white-noise approach. Springer Science & Business Media, 2012.

6. Melinda E Koelling and Duane Q Nykamp. Computing linear approximations to non-linear neuronal response. Network: Computation in Neural Systems, 19(4):286–313, 2008.

7. Tim Gollisch and Markus Meister. Eye smarter than scientists believed: neural computations in circuits of the retina. Neuron, 65(2):150–164, 2010.

8. Esteban Real, Hiroki Asari, Tim Gollisch, and Markus Meister. Neural circuit inference from function to structure. Current Biology, 27(2):189–198, 2017.

9. Ben Willmore, Ryan J Prenger, Michael C-K Wu, and Jack L Gallant. The berkeley wavelet transform: a biologically inspired orthogonal wavelet transform. Neural computation, 20(6):1537–1564, 2008.

10. Niru Maheswaranathan, David B Kastner, Stephen A Baccus, and Surya Ganguli. Inferring hidden structure in multilayered neural circuits. PLoS computational biology, 14 (8):e1006291, 2018.

11. Daniel LK Yamins, Ha Hong, Charles F Cadieu, Ethan A Solomon, Darren Seibert, and James J DiCarlo. Performance-optimized hierarchical models predict neural responses in higher visual cortex. Proceedings of the National Academy of Sciences, 111(23): 8619–8624, 2014.

12. Umut Güçlü and Marcel AJ van Gerven. Deep neural networks reveal a gradient in the complexity of neural representations across the ventral stream. Journal of Neuroscience, 35(27):10005–10014, 2015.

13. Yann LeCun, Yoshua Bengio, and Geoffrey Hinton. Deep learning. nature, 521(7553): 436–444, 2015.

14. Demis Hassabis, Dharshan Kumaran, Christopher Summerfield, and Matthew Botvinick. Neuroscience-inspired artificial intelligence. Neuron, 95(2):245–258, 2017.

15. Maxwell H Turner, Luis Gonzalo Sanchez Giraldo, Odelia Schwartz, and Fred Rieke. Stimulus-and goal-oriented frameworks for understanding natural vision. Nature neuroscience, 22(1):15–24, 2019.

16. Blake A Richards, Timothy P Lillicrap, Philippe Beaudoin, Yoshua Bengio, Rafal Bogacz, Amelia Christensen, Claudia Clopath, Rui Ponte Costa, Archy de Berker, Surya Ganguli, et al. A deep learning framework for neuroscience. Nature neuroscience, 22(11):1761–1770, 2019.

17. Daniel LK Yamins and James J DiCarlo. Using goal-driven deep learning models to understand sensory cortex. Nature neuroscience, 19(3):356–365, 2016.

18. Lane McIntosh, Niru Maheswaranathan, Aran Nayebi, Surya Ganguli, and Stephen Baccus. Deep learning models of the retinal response to natural scenes. Advances in neural information processing systems, 29:1369–1377, 2016.

19. David Klindt, Alexander S Ecker, Thomas Euler, and Matthias Bethge. Neural system identification for large populations separating “what” and “where”. in Advances in Neural Information Processing Systems, pages 3506–3516, 2017.

20. Pouya Bashivan, Kohitij Kar, and James J DiCarlo. Neural population control via deep image synthesis. Science, 364(6439), 2019.

21. Carlos R Ponce, Will Xiao, Peter F Schade, Till S Hartmann, Gabriel Kreiman, and Margaret S Livingstone. Evolving images for visual neurons using a deep generative network reveals coding principles and neuronal preferences. Cell, 177(4):999–1009, 2019.

22. Edgar Y Walker, Fabian H Sinz, Erick Cobos, Taliah Muhammad, Emmanouil Froudarakis, Paul G Fahey, Alexander S Ecker, Jacob Reimer, Xaq Pitkow, and Andreas S Tolias. Inception loops discover what excites neurons most using deep predictive models. Nature neuroscience, 22(12):2060–2065, 2019.

23. Tom Baden, Thomas Euler, and Philipp Berens. Understanding the retinal basis of vision across species. Nature Reviews Neuroscience, 21(1):5–20, 2020.

24. Horace B Barlow et al. Possible principles underlying the transformation of sensory messages. Sensory communication, 1(01), 1961.

25. Eero P Simoncelli and Bruno A Olshausen. Natural image statistics and neural representation. Annual review of neuroscience, 24(1):1193–1216, 2001.

26. Eugene Switkes, Melanie J Mayer, and Jeffrey A Sloan. Spatial frequency analysis of the visual environment: Anisotropy and the carpentered environment hypothesis. Vision research, 18(10):1393–1399, 1978.

27. Xiangmin Xu, Christine E Collins, Ilya Khaytin, Jon H Kaas, and Vivien A Casagrande. Unequal representation of cardinal vs. oblique orientations in the middle temporal visual area. Proceedings of the National Academy of Sciences, 103(46):17490–17495, 2006.

28. Ahna R Girshick, Michael S Landy, and Eero P Simoncelli. Cardinal rules: visual orientation perception reflects knowledge of environmental statistics. Nature neuroscience, 14(7):926–932, 2011.

29. Simon Laughlin. A simple coding procedure enhances a neuron’s information capacity. Zeitschrift für Naturforschung c, 36(9-10):910–912, 1981.

30. J Hans van Hateren and Dan L Ruderman. Independent component analysis of natural image sequences yields spatio-temporal filters similar to simple cells in primary visual cortex. Proceedings of the Royal Society of London. Series B: Biological Sciences, 265 (1412):2315–2320, 1998.

31. Suva Roy, Na Young Jun, Emily L Davis, John Pearson, and Greg D Field. Inter-mosaic coordination of retinal receptive fields. Nature, 592(7854):409–413, 2021.

32. Joseph J Atick and A Norman Redlich. Towards a theory of early visual processing. Neural computation, 2(3):308–320, 1990.

33. Joseph J Atick. Could information theory provide an ecological theory of sensory processing? Network: Computation in neural systems, 3(2):213–251, 1992.

34. Christina Enroth-Cugell and John G Robson. The contrast sensitivity of retinal ganglion cells of the cat. The Journal of physiology, 187(3):517–552, 1966.

35. Dana H Ballard. Modular learning in neural networks. in AAAI, pages 279–284, 1987.

36. Geoffrey E Hinton and Ruslan R Salakhutdinov. Reducing the dimensionality of data with neural networks. science, 313(5786):504–507, 2006.

37. Samuel Ocko, Jack Lindsey, Surya Ganguli, and Stephane Deny. The emergence of multiple retinal cell types through efficient coding of natural movies. in Advances in Neural Information Processing Systems, pages 9389–9400, 2018.

38. Yongrong Qiu, Zhijian Zhao, David Klindt, Magdalena Kautzky, Klaudia P Szatko, Frank Schaeffel, Katharina Rifai, Katrin Franke, Laura Busse, and Thomas Euler. Natural environment statistics in the upper and lower visual field are reflected in mouse retinal specializations. Current Biology, 2021.

39. Dylan M Paiton, Charles G Frye, Sheng Y Lundquist, Joel D Bowen, Ryan Zarcone, and Bruno A Olshausen. Selectivity and robustness of sparse coding networks. Journal of Vision, 20(12):10–10, 2020.

40. Jan Eichhorn, Fabian Sinz, and Matthias Bethge. Natural image coding in v1: how much use is orientation selectivity? PLoS computational biology, 5(4):e1000336, 2009.

41. Wiktor Mlynarski, Michal Hledík, Thomas R Sokolowski, and Gašper Tkačik. Statistical analysis and optimality of neural systems. Neuron, 109(7):1227–1241, 2021.

42. Benjamin T Vincent and Roland J Baddeley. Synaptic energy efficiency in retinal processing. Vision research, 43(11):1285–1292, 2003.

43. Tom Baden, Philipp Berens, Katrin Franke, Miroslav Román Rosón, Matthias Bethge, and Thomas Euler. The functional diversity of retinal ganglion cells in the mouse. Nature, 529(7586):345–350, 2016.

44. Klaudia P Szatko, Maria M Korympidou, Yanli Ran, Philipp Berens, Deniz Dalkara, Timm Schubert, Thomas Euler, and Katrin Franke. Neural circuits in the mouse retina support color vision in the upper visual field. Nature communications, 11(1):1–14, 2020.

45. Cassandra L Schlamp, Angela D Montgomery, Caitlin E Mac Nair, Claudia Schuart, Daniel J Willmer, and Robert W Nickells. Evaluation of the percentage of ganglion cells in the ganglion cell layer of the rodent retina. Molecular vision, 19:1387, 2013.

46. Gerald H Jacobs, Gary A Williams, and John A Fenwick. Influence of cone pigment coexpression on spectral sensitivity and color vision in the mouse. Vision research, 44 (14):1615–1622, 2004.

47. Matthew D Zeiler and Rob Fergus. Visualizing and understanding convolutional networks. in European conference on computer vision, pages 818–833. Springer, 2014.

48. Katrin Franke, Philipp Berens, Timm Schubert, Matthias Bethge, Thomas Euler, and Tom Baden. Inhibition decorrelates visual feature representations in the inner retina. Nature, 542(7642):439–444, 2017.

49. Robert E Soodak. Two-dimensional modeling of visual receptive fields using gaussian subunits. Proceedings of the National Academy of Sciences, 83(23):9259–9263, 1986.

50. Rajesh PN Rao and Dana H Ballard. Predictive coding in the visual cortex: a functional interpretation of some extra-classical receptive-field effects. Nature neuroscience, 2(1): 79–87, 1999.

51. Toshihiko Hosoya, Stephen A Baccus, and Markus Meister. Dynamic predictive coding by the retina. Nature, 436(7047):71–77, 2005.

52. Jamie Johnston, Sofie-Helene Seibel, Léa Simone Adele Darnet, Sabine Renninger, Michael Orger, and Leon Lagnado. A retinal circuit generating a dynamic predictive code for oriented features. Neuron, 102(6):1211–1222, 2019.

53. J Hans van Hateren. Real and optimal neural images in early vision. Nature, 360(6399): 68–70, 1992.

54. Joseph J Atick and A Norman Redlich. What does the retina know about natural scenes? Neural computation, 4(2):196–210, 1992.

55. Matthew Chalk, Olivier Marre, and Gašper Tkačik. Toward a unified theory of efficient, predictive, and sparse coding. Proceedings of the National Academy of Sciences, 115 (1):186–191, 2018.

56. Jerome Y Lettvin, Humberto R Maturana, Warren S McCulloch, and Walter H Pitts. What the frog’s eye tells the frog’s brain. Proceedings of the IRE, 47(11):1940–1951, 1959.

57. J Alexander Bae, Shang Mu, Jinseop S Kim, Nicholas L Turner, Ignacio Tartavull, Nico Kemnitz, Chris S Jordan, Alex D Norton, William M Silversmith, Rachel Prentki, et al. Digital museum of retinal ganglion cells with dense anatomy and physiology. Cell, 173 (5):1293–1306, 2018.

58. Nicholas M Tran, Karthik Shekhar, Irene E Whitney, Anne Jacobi, Inbal Benhar, Guosong Hong, Wenjun Yan, Xian Adiconis, McKinzie E Arnold, Jung Min Lee, et al. Single-cell profiles of retinal ganglion cells differing in resilience to injury reveal neuroprotective genes. Neuron, 104(6):1039–1055, 2019.

59. Jillian Goetz, Zachary F Jessen, Anne Jacobi, Adam Mani, Sam Cooler, Devon Greer, Sabah Kadri, Jeremy Segal, Karthik Shekhar, Joshua Sanes, et al. Unified classification of mouse retinal ganglion cells using function, morphology, and gene expression. Morphology, and Gene Expression, 2021.

60. Horace B Barlow and Richard M Hill. Selective sensitivity to direction of movement in ganglion cells of the rabbit retina. Science, 139(3553):412–412, 1963.

61. Fabian H Sinz, Alexander S Ecker, Paul G Fahey, Edgar Y Walker, Erick Cobos, Emmanouil Froudarakis, Dimitri Yatsenko, Xaq Pitkow, Jacob Reimer, and Andreas S Tolias. Stimulus domain transfer in recurrent models for large scale cortical population prediction on video. BioRxiv, page 452672, 2018.

62. Jon Touryan, Gidon Felsen, and Yang Dan. Spatial structure of complex cell receptive fields measured with natural images. Neuron, 45(5):781–791, 2005.

63. Andrew Saxe, Stephanie Nelli, and Christopher Summerfield. If deep learning is the answer, what is the question? Nature Reviews Neuroscience, 22(1):55–67, 2021.

64. Alexander Heitman, Nora Brackbill, Martin Greschner, Alexander Sher, Alan M Litke, and EJ Chichilnisky. Testing pseudo-linear models of responses to natural scenes in primate retina. bioRxiv, page 045336, 2016.

65. Nicole C Rust and J Anthony Movshon. In praise of artifice. Nature neuroscience, 8 (12):1647–1650, 2005.

66. Yifeng Zhang, In-Jung Kim, Joshua R Sanes, and Markus Meister. The most numerous ganglion cell type of the mouse retina is a selective feature detector. Proceedings of the National Academy of Sciences, 109(36):E2391–E2398, 2012.

67. Horace B Barlow. Summation and inhibition in the frog’s retina. The Journal of physiology, 119(1):69–88, 1953.

68. Sophie Deneve and Matthew Chalk. Efficiency turns the table on neural encoding, decoding and noise. Current Opinion in Neurobiology, 37:141–148, 2016.

69. Michael Teti, Emily Meyer, and Garrett Kenyon. Can lateral inhibition for sparse coding help explain v1 neuronal responses to natural stimuli? In 2020 IEEE Southwest Symposium on Image Analysis and Interpretation (SSIAI), pages 120–124. IEEE, 2020.

70. Horace Barlow. Redundancy reduction revisited. Network: computation in neural systems, 12(3):241, 2001.

71. Brett Vintch, J Anthony Movshon, and Eero P Simoncelli. A convolutional subunit model for neuronal responses in macaque v1. Journal of Neuroscience, 35(44):14829–14841, 2015.

72. Santiago A Cadena, George H Denfield, Edgar Y Walker, Leon A Gatys, Andreas S Tolias, Matthias Bethge, and Alexander S Ecker. Deep convolutional models improve predictions of macaque v1 responses to natural images. PLoS computational biology, 15(4):e1006897, 2019.

73. Kevin L Briggman and Thomas Euler. Bulk electroporation and population calcium imaging in the adult mammalian retina. Journal of neurophysiology, 105(5):2601–2609, 2011.

74. Thomas Euler, Susanne E Hausselt, David J Margolis, Tobias Breuninger, Xavier Castell, Peter B Detwiler, and Winfried Denk. Eyecup scope—optical recordings of light stimulus-evoked fluorescence signals in the retina. Pflügers Archiv-European Journal of Physiology, 457(6):1393–1414, 2009.

75. Thomas Euler, Katrin Franke, and Tom Baden. Studying a light sensor with light: multiphoton imaging in the retina. in Multiphoton Microscopy, pages 225–250. Springer, 2019.

76. Katrin Franke, André Maia Chagas, Zhijian Zhao, Maxime JY Zimmermann, Philipp Bartel, Yongrong Qiu, Klaudia P Szatko, Tom Baden, and Thomas Euler. An arbitrary-spectrum spatial visual stimulator for vision research. elife, 8:e48779, 2019.

77. Leo Breiman. Random forests. Machine learning, 45(1):5–32, 2001.

78. F. Pedregosa, G. Varoquaux, A. Gramfort, V. Michel, B. Thirion, O. Grisel, M. Blondel, P. Prettenhofer, R. Weiss, V. Dubourg, J. Vanderplas, A. Passos, D. Cournapeau, M. Brucher, M. Perrot, and E. Duchesnay. Scikit-learn: Machine learning in Python. Journal of Machine Learning Research, 12:2825–2830, 2011.

79. Lin Sun, Kui Jia, Dit-Yan Yeung, and Bertram E Shi. Human action recognition using factorized spatio-temporal convolutional networks. in Proceedings of the IEEE international conference on computer vision, pages 4597–4605, 2015.

80. Du Tran, Heng Wang, Lorenzo Torresani, Jamie Ray, Yann LeCun, and Manohar Paluri. A closer look at spatiotemporal convolutions for action recognition. in Proceedings of the IEEE conference on Computer Vision and Pattern Recognition, pages 6450–6459, 2018.

81. Eizaburo Doi and Michael S Lewicki. A theory of retinal population coding. Advances in neural information processing systems, 19:353, 2007.

82. MCW Van Rossum, Brendan J O’Brien, and Robert G Smith. Effects of noise on the spike timing precision of retinal ganglion cells. Journal of neurophysiology, 89(5):2406–2419, 2003.

83. David J Field. What is the goal of sensory coding? Neural computation, 6(4):559–601, 1994.

84. David H Hubel and Torsten N Wiesel. Receptive fields of single neurones in the cat’s striate cortex. The Journal of physiology, 148(3):574–591, 1959.

85. David Marr and Ellen Hildreth. Theory of edge detection. Proceedings of the Royal Society of London. Series B. Biological Sciences, 207(1167):187–217, 1980.

